# Lysine acetylation-mediated regulation of ferredoxin and ferredoxin reductase redox-active proteins in *Haloferax volcanii*

**DOI:** 10.64898/2026.08.10.743930

**Authors:** Katherine R. Weber, Allie Aguila, Semaj Bulter-Drinks, Peter Huynh, Brianna Novillo, Xin Wang, Christian Heryakusuma, Biswarup Mukhopadhyay, Julie A. Maupin-Furlow

## Abstract

Lysine acetylation is an evolutionarily conserved, post-translational modification that regulates metabolism and protein function, yet its role in archaeal electron transfer systems remains poorly understood. Here, we investigated lysine acetylation of the 2Fe–2S ferredoxin *Hv*Fdx (HVO_2995) and its flavin-dependent oxidoreductase *Hv*FdR (HVO_2345) partner in the halophilic archaeon *Haloferax volcanii*. Genetic and biochemical analyses established *Hv*Fdx as an essential 2Fe–2S ferredoxin with a midpoint redox potential of −385 mV. Lysine acetylation of *Hv*Fdx was found to occur primarily at K119, a residue positioned near the [Fe-S] cluster interface, and to modulate electron transfer capacity without impacting Fe–S cluster incorporation, midpoint potential, or protein abundance. In contrast, *Hv*FdR was found lysine acetylated at multiple sites in a manner consistent with a non-enzymatic mechanism that resulted in altered flavin binding, enzymatic activity, and thermal stability. Lysine acetylation of *Hv*Fdx was found to stimulate electron flow from *Hv*FdR as measured by an anaerobic NADPH → *Hv*FdR → *Hv*Fdx → DCIP assay. 3D structural modeling, proteomic, biochemical, and genetic assays suggest the haloarchaeal GNAT-family acetyltransferase homolog HVO_2874 as a candidate enzyme associated with *Hv*Fdx lysine acetylation and optimal growth of *H. volcanii*. Together, these findings demonstrate that lysine acetylation differentially regulates archaeal redox-active proteins and functions as an important mechanism coordinating redox metabolism in *H. volcanii*.

## Introduction

Haloarchaea inhabit hypersaline environments characterized by extreme conditions including ultraviolet (UV) irradiation [1], desiccation and rehydration (hypo/hyper-osmolarity) [2], extreme temperatures and pH [3, 4], heavy metals [5], and chloride or other oxidative agents [6]. Survival under these conditions requires tightly coordinated metabolic and redox systems capable of maintaining cellular redox homeostasis and protecting against oxidative damage [7–10]. Central to these processes are ferredoxins and flavin-based ferredoxin reductases, which function as electron carriers and mediators, respectively, that shuttle electrons between metabolic pathways and contribute to intracellular redox balance [11, 12]. These redox-active proteins support diverse cellular processes such as photosynthesis [13], nitrogen metabolism [14], hydrogen generation [15], ethanol production [16], and [Fe-S] cluster assembly [17]. Despite their cellular importance, the regulatory mechanisms governing archaeal redox-active proteins remain poorly understood.

Post-translational modifications (PTMs), particularly lysine acetylation, have emerged as important regulators of protein function across all domains of life [18–22]. Lysine acetylation can alter protein charge, conformation, cofactor binding, stability, protein interactions, and catalytic activity, thereby enabling rapid adaptation of cellular processes in response to environmental change [20, 23, 24]. Although initially characterized as a regulator of eukaryotic histones, lysine acetylation is now recognized as an evolutionarily conserved mechanism that regulates metabolism, stress adaptation and redox homeostasis in both prokaryotes and eukaryotes [25–29].

Lysine acetylation is suggested to be a widespread mechanism for regulating electron transfer systems across diverse organisms. Early studies reveal a 2Fe–2S ferredoxin (Fdx) that is acetylated on a conserved lysine in the halophilic archaea *Halobacterium salinarum* [30] and *Haloarcula marismortui* [31], providing the first evidence of lysine acetylation in prokaryotes and demonstrating that lysine acetylation can occur on redox-active proteins. Subsequent acetylome studies reveal this 2Fe–2S Fdx is similarly acetylated at the same conserved lysine 119 in *Haloferax mediterranei* [32] and *Haloferax volcanii* [6], suggesting an evolutionarily conserved regulatory mechanism linked to ferredoxin-dependent electron transfer. In parallel, lysine acetylation of ferredoxin-dependent oxidoreductases is identified across diverse biological systems including the flavin-dependent oxidoreductase homologs of *H. volcanii* (*Hv*FdR, HVO_2345) [6] and *Hx. mediterranei* (HFX_2354) [32], as well as plant ferredoxin–NADP^+^ oxidoreductases, such as that from *Arabidopsis thaliana*, where lysine residues near the NADP(H)-binding region are modified [33]. These findings suggest a conserved role for lysine acetylation in modulating co-factor interactions and redox activity. Large-scale acetylome studies demonstrate oxidoreductases and electron carrier proteins are frequently lysine acetylated, suggesting an important role for this PTM in controlling redox metabolism and electron transfer pathways [34–37]. Collectively, these findings support the idea that lysine acetylation is a widespread mechanism for regulating electron transfer systems across diverse organisms.

Although lysine acetylation of haloarchaeal ferredoxins has been recognized for more than forty years, its functional significance and the enzymes responsible for establishing these modifications have remained largely unknown. In this study, the role of lysine acetylation in regulating electron transfer was investigated with the [2Fe-2S] ferredoxin *Hv*Fdx (FerA5; HVO_2995) and its redox partner, the flavin-dependent oxidoreductase *Hv*FdR (HVO_2345), from *H. volcanii*. Genetic analyses demonstrated that *fdx* is essential for viability, consistent with the reduced growth phenotype previously observed in an *fdr* mutant (Weber *et al*., submitted), highlighting the central importance of this electron-transfer pathway to cellular physiology. Biochemical, structural, genetic, and proteomic analyses identified K119 as the primary site of lysine acetylation on *Hv*Fdx and demonstrated that *Hv*FdR is acetylated at multiple lysine residues. Functional characterization revealed that *Hv*Fdx acetylation influences electron transfer within the *Hv*FdR–*Hv*Fdx pathway while exerting minimal effects on the intrinsic properties of *Hv*Fdx, including Fe–S cluster assembly, midpoint redox potential, and protein abundance. These findings support a model in which lysine acetylation regulates electron flux primarily by altering electron transfer efficiency and interactions with redox partners. Further investigation identified HVO_2874 as a putative GNAT-family acetyltransferase associated with *Hv*Fdx acetylation, suggesting a role in the modification of this highly conserved haloarchaeal ferredoxin. In contrast, lysine acetylation of *Hv*FdR appears to occur predominantly through a non-enzymatic mechanism and influences flavin binding, catalytic activity, and thermal stability. Collectively, these findings resolve a longstanding question regarding the functional significance and regulation of ferredoxin acetylation and establish lysine acetylation as a key regulatory mechanism governing archaeal electron transfer and redox homeostasis.

## Results

### *Hv*Fdx plays an important role in *H. volcanii* physiology

To investigate the physiological role of *Hv*Fdx and lysine acetylation, genetic approaches were first employed. However, attempts to generate an *fdx* (*ferA5*) deletion strain through homologous recombination [38] or conditional depletion [39] were unsuccessful. A CRISPR interference (CRISPRi)-based suppression strategy [40, 41] was therefore used to reduce *fdx* (*ferA5*) expression without complete disruption of the gene. CRISPR RNAs (crRNAs) were designed to target the promoter region and template strand of *fdx*, as these regions have previously been shown to yield strong transcriptional repression of other operons in *H. volcanii* [40]. Four independent CRISPRi constructs (CP1–CP4, CRISPRi plasmids) targeting distinct regions of the *fdx* (*ferA5*) template strand were generated and expressed in the CRISPRi host strain HV30 (Δ*cas3*Δ*bgaH*Δ*cas6b*) (**Fig. 1A**). To evaluate repression efficiency, *fdx* transcript abundance was quantified by qRT–PCR relative to the HV30/EV (empty vector) control. Compared to HV30/EV, CRISPRi constructs CP1, CP2, CP3, and CP4 reduced *fdx* transcript abundance by 38%, 73%, 79%, and 24%, respectively (**Fig. 1B**), demonstrating effective and variable suppression across the targeted regions.

**Figure 1.**
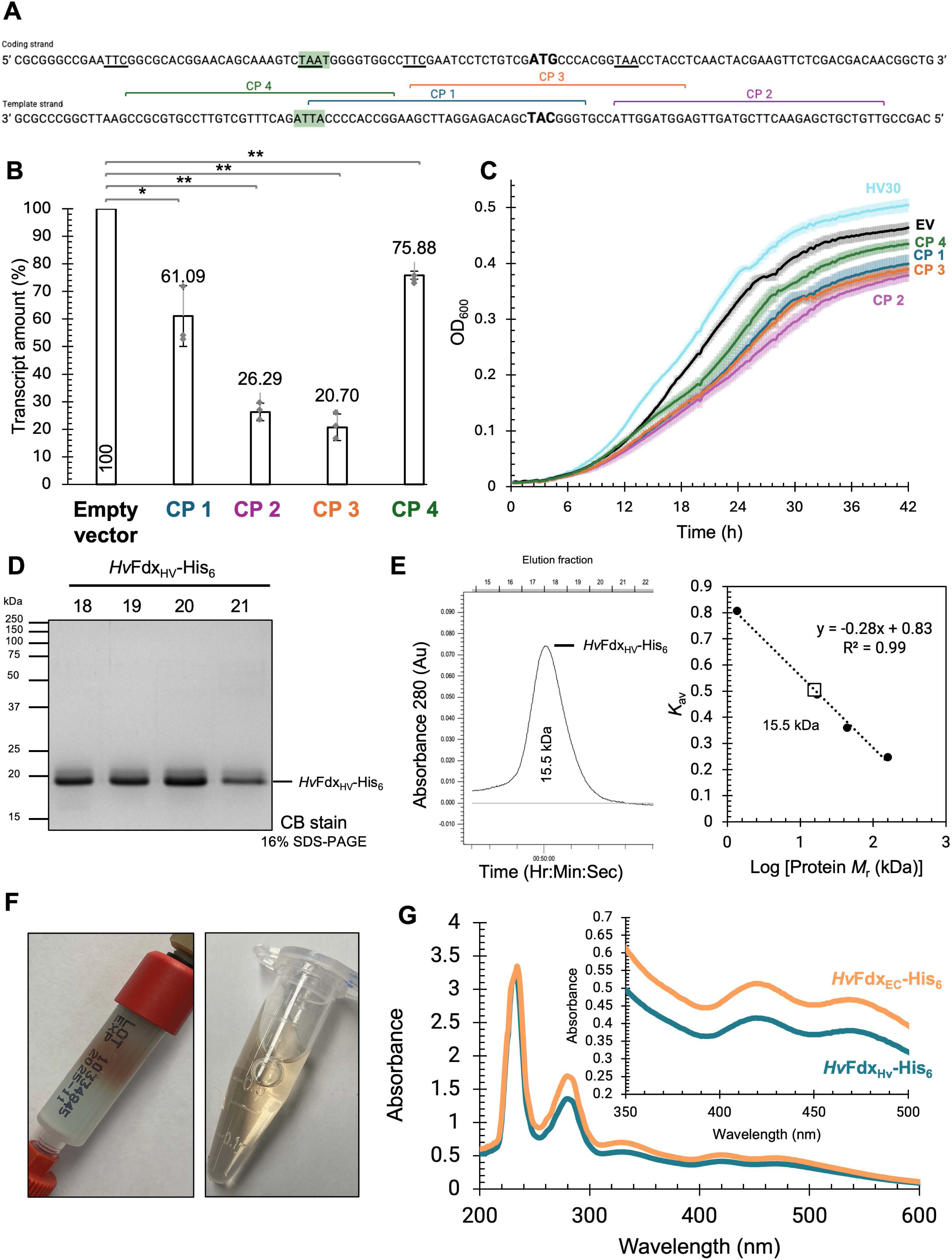
CRISPRi-mediated repression and biochemical characterization of *Hv*Fdx. A. Schematic representation of CRISPR interference (CRISPRi) guide RNA targets the template strand within the *fdx* promoter and coding region. The coding and template DNA strands are shown in the 5′→3′ and 3′→5′ orientations, respectively. The predicted promoter TATA box is highlighted in green, and the translational start codon (ATG/TAC) is shown in bold. Guide RNA target regions are indicated by colored brackets corresponding to individual CRISPRi constructs: CP1 (blue), CP2 (purple), CP3 (orange), and CP4 (green). Guides were designed to target either the promoter region or early coding sequence to suppress *fdx* transcription. B. Effect of CRISPRi-mediated repression on *fdx* transcript abundance determined by qRT-PCR. Relative fold changes in *fdx* transcript levels were quantified using the ΔΔCt method and normalized to the internal reference gene *rpL-16*. Transcript abundance was compared between CRISPRi-targeted strains and the non-targeting control. Data shown as mean ± SD (*n* = 3 biological replicates and 3 technical replicates). Strip boxes, log phase; solid boxes, stationary phase. Student’s *t*-test used to determine statistical significance (*p*-value ≤ 0.005, **; ≤ 0.05, *). HV30/EV verses HV30/pJAM4489 (*p* = 0.02), HV30/pJAM4490 (*p* = 0.0005), HV30/pJAM4491 (*p* = 0.001), and HV30/pJAM4492 (*p* = 0.0008). C. *H. volcanii* HV30 CRISPRi impairs growth. *H. volcanii* strains included: HV30 (parent, light blue), HV30/pJAM4484 (empty vector, black), HV30/pJAM4489 (CRISPRi plasmid *fdx* 1, dark blue), HV30/pJAM4490 (CRISPRi *fdx* plasmid 2, purple), HV30/pJAM4491 (CRISPRi *fdx* plasmid 3, orange), HV30/ pJAM4491 (CRISPRi *fdx* plasmid 4, green). Growth curve at 42 °C in ATCC974 rich medium supplemented with novobiocin for plasmid selection. Growth was then monitored by OD_600_ using EPOCH2 microplate reader and Gen5 software, every 15 min for 99 h, shaking at double orbital (continuously) (see Methods for details). D. H1207/pJAM4455 was used to express *Hv*Fdx with a C-terminal His_6_-tag using an inducible *ptnA* promoter. Cells were grown to log phase (OD_600_ 0.6-0.8) on Hv-Ca^+^ medium, induced with 2 mM tryptophan, and grown to stationary phase (OD_600_ 1.2). *Hv*Fdx-His_6_ was purified by Ni-NTA affinity (HisTrap) and Superdex 75 10/300 GL (SEC) chromatography (see Methods for details). *Hv*Fdx-His_6_ purifies to apparent homogeneity based on analysis of SEC fractions 18 to 21 (1 µg per lane) by reducing 16% SDS-PAGE and Coomassie blue staining (CB stain). E. *Hv*Fdx-His_6_ purifies as a monomer based on SEC. Left: SEC chromatogram where fraction 14.9 correlates to an observed molecular weight of 15.5 kDa. Right: SEC standard curve. His_6_-*Hv*FdR (open square) displays an observed molecular weight of 15.5 kDa, with a theoretical molecular weight of 15.5 kDa, compared to the standards (closed circles). F. *Hv*Fdx-His_6_ likely binds iron based on the brown coloration observed during HisTrap chromatography (left) and after elution from the column (right). G. *Hv*Fdx-His_6_ contains bound [2Fe-2S] based on UV-Vis spectrum and iron-sulfur content assays of purified *Hv*Fdx-His_6_ expressed in *H. volcanii* and *E. coli*. X-axis displays absorbance and y-axis wavelength. When *Hv*Fdx-His_6_ is expressed in both species, it displays two maximum peaks at 418 and 468 nm (see Methods for details).

To determine the physiological consequences of *fdx* (*ferA5*, *Hv*Fdx) repression, growth of the CRISPRi strains was evaluated in rich medium. Relative to the HV30/EV control, all crRNAs exhibited impaired growth characterized by reduced area under the growth curve (AUC) and decreased max optical density 600 (OD_600_), as well as an increased doubling time (**Fig. 1C, Supplemental Fig. S1A-B**). The most pronounced growth defect was observed with the HV30 strain carrying the CRISPR *fdx* plasmid 2 (CP2), which displayed the second strongest reduction in *fdx* transcript abundance by qRT–PCR, followed closely by CRISPR *fdx* plasmid 3 (CP3) that displayed the strongest reduction in *fdx* transcript. Overall, a direct relationship between *fdx* transcript suppression and impaired cellular fitness was observed. Collectively, these findings indicate that expression of *fdx* is required for optimal growth and strongly support an essential role for *Hv*Fdx in *H. volcanii* physiology. Notably, while the gene (*fdr*, *hvo_2345*) encoding the FAD-dependent oxidoreductase *Hv*FdR that can partner with *Hv*Fdx in electron transfer is nonessential, deletion of *fdr* results in a pronounced reduction in growth relative to the wild-type (Weber *et al*., submitted). This difference in essentiality suggests that *Hv*Fdx likely participates in multiple electron transfer pathways and interacts with additional redox partners beyond *Hv*FdR, consistent with its central role as a versatile electron carrier in cellular metabolism. Together, these results suggest that *Hv*Fdx contributes importantly to the cellular fitness of *H. volcanii*.

### Purification of *Hv*Fdx as a soluble monomer

To characterize the biochemical properties of *Hv*Fdx, the encoding gene *hvo_2995* was cloned into a modified inducible\ *H. volcanii* expression vector (pTA963) to generate *Hv*Fdx_HV_-His_6_ and transformed into *H. volcanii* H1207. The construct was designed to replace the N-terminal His-tag encoded by pTA963 with a C-terminal His-tag, as N-terminal tag (constitutive or inducible) did not yield sufficient *Hv*Fdx [2Fe-2S] protein for biochemical study. Cells were cultivated in Hv-Ca^+^ medium, supplemented with 2 mM ammonium iron (III) citrate and 2 mM L-cysteine, and protein expression was induced with 2 mM tryptophan. *Hv*Fdx-His_6_ was subsequently purified by Ni-NTA affinity chromatography followed by size exclusion chromatography. SDS-PAGE (16%) analysis of the SEC-purified *Hv*Fdx-His_6_ revealed a predominant protein band migrating at approximately 20 kDa (**Fig. 1D**). The observed doublet band is proposed to result from differences in the redox state or from conformational changes associated with protein unfolding and disulfide bond formation, as observed with related proteins [42–44]. SEC analysis demonstrated that *Hv*Fdx-His_6_ eluted as a single symmetrical peak (**Fig. 1E, left**), corresponding to an observed molecular mass of 15.5 kDa (**Fig. 1E, right**). These findings indicate that *Hv*Fdx-His_6_ adopts a monomeric conformation characteristic of previously characterized 2Fe–2S ferredoxins [45, 46]. Collectively, these results demonstrate that *Hv*Fdx-His_6_ can be purified as a soluble, monomeric protein.

### *Hv*Fdx contains a [2Fe-2S] cluster as its cofactor

*Hv*Fdx was predicted to coordinate an iron-sulfur cluster based on characterized [2Fe-2S] ferredoxin homologs [47–50]. The purified *Hv*Fdx-His_6_ displayed a distinct brown coloration (**Fig. 1F**), consistent with [Fe-S] cluster incorporation. To evaluate [Fe-S] binding, the *Hv*Fdx-His_6_ protein samples expressed and purified from *H. volcanii* (*Hv*Fdx_HV_-His_6_) and *E. coli* (*Hv*Fdx_EC_-His_6_) were compared by UV–visible spectroscopy under aerobic conditions. Both protein preparations exhibited maxima at A_418_ and A_468_ (**Fig. 1G**), spectral features of known oxidized [2Fe–2S] ferredoxins [51, 52], thus supporting incorporation of the Fe–S cluster in the native host (*H. volcanii*) and bacterial expression system (*E. coli*). Iron quantification further confirmed Fe incorporation into *Hv*Fdx-His_6_ purified from *H. volcanii* (Fe, 2.09 ± 0.07 µM) and *E. coli* (Fe, 2.04 ± 0.08 µM). Together, these findings demonstrate that a [2Fe-2S] cluster is coordinated in the *Hv*Fdx-His_6_ protein independent of the expression host, enabling downstream biochemical and redox characterization.

### 3D structural modeling of *Hv*Fdx identifies lysine acetylation site K119 proximal to the [Fe–S] cluster

Previous SILAC-based lysine acetylome profiling identified *Hv*Fdx to be one of the most highly lysine acetylated proteins in the *H. volcanii* proteome and to be modified at multiple sites, including K97, K113 and K119 [6]. Multiple sequence alignment reveals the 2Fe–2S cluster-coordinating cysteine residues and the K119 site are highly conserved among haloarchaeal [2Fe-2S] ferredoxin homologs (**Fig. 2A, Supplemental Table S1**). AlphaFold 3D-modeling suggests all three sites of lysine acetylation are exposed on the *Hv*Fdx surface, with K119 located proximal to the [2Fe–2S] cluster-binding region (**Fig. 2B and 2C**). Coulombic surface electrostatic analysis further suggests the sites of lysine acetylation are located within acidic regions of net negative electrostatic charge on the surface. The predicted spatial proximity of K119 to the [Fe–S] cluster (**Fig. 2C**), suggests lysine acetylation at this site could influence the electron transfer properties of *Hv*Fdx. Collectively, these findings suggest a conserved and potentially important regulatory role for lysine acetylation in haloarchaeal [2Fe-2S] ferredoxin function.

**Figure 2.**
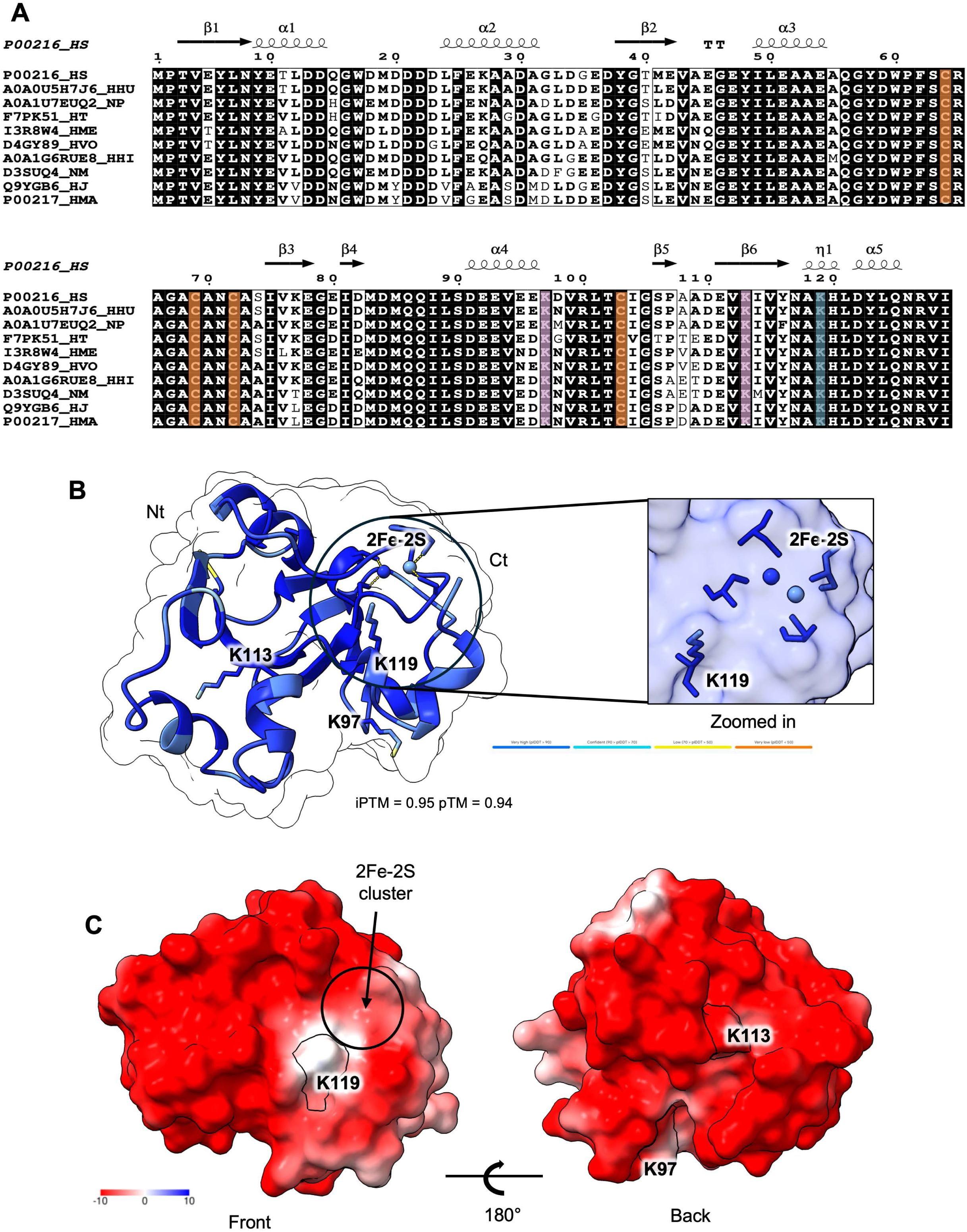
Structural features of *Hv*Fdx (FerA5 [2Fe-2S] ferredoxin, HVO_2995) acetylation. A. Acetylated lysine K119 is conserved in other of halophilic [2Fe-2S] ferredoxins. The alignment highlights conserved features among archaeal halophilic [2Fe-2S] ferredoxins homologs. All four cysteine residues involved in [2Fe-2S] cluster coordination are conserved across all species (orange). Conserved lysine residues at positions 113 and 119 are conserved (purple). Lysine 119 (green) is found to be acetylated in *Haloarcula marismortui*, *Halobacterium salinarum, Haloferax mediterranei*, and *Haloferax volcanii*, suggesting a potential regulatory or structural role. Lysine 77 is not conserved in *Natrialba magadii*, *Haloarcula japonica*, and *Haloarcula marismortui* (blue). Uniprot accession numbers are provided. Abbreviations: HS, *Halobacterium salinarum*; HHU, *Halobacterium hubeiense*; NP, Natronomonas pharaonic; HT, *Halorhabdus tiamatea*; HME, *Haloferax mediterranei*; HVO, *Haloferax volcanii*; HHI, *Haloterrigena hispanica*; NM, *Natrialba magadii*; HJ, *Haloarcula japonica*; HMA, *Haloarcula marismortui*. Multialign was used to generate the sequence alignment and imported to ESPript 3.0. *Halobacterium salinarum* PDBe ID 1e10 was used as the input file for the secondary structure depiction. B. AlphaFold server predicted model (ipTM = 0.95 pTM = 0.94) with *Hv*Fdx from *H. volcanii* with 2 Fe^3+^ ions. (Left) Model of *Hv*Fdx using AlphaFold server, with KAc sites K97, K113, and K119. (Right) Predicted [Fe-S] cluster and K119 are in close proximity. C. Surface charge models of *Hv*Fdx. Positive (blue) and neutral (white) charges are predicted to be protein-protein interaction sites.

### *Hv*Fdx is lysine acetylated in *H. volcanii* but not in recombinant *E. coli*

To assess the lysine acetylation status of *Hv*Fdx-His_6_ purified from *H. volcanii* and recombinant *E. coli*, the proteins were analyzed by anti-acetyllysine immunoblotting along with Coomassie blue staining and anti-His immunoblotting (**Fig. 3A**). The *Hv*Fdx-His_6_ expressed in *H. volcanii* was found to be lysine acetylated, while the protein expressed in recombinant *E. coli* lacked a detectable lysine acetylation signal. These results reveal that *Hv*Fdx is acetylated in its native host but not in recombinant *E. coli*, providing two distinct lysine acetylated states of the protein for further characterization. This suggests the modification may be regulated by an enzymatic mechanism specific to the native host.

**Figure 3.**
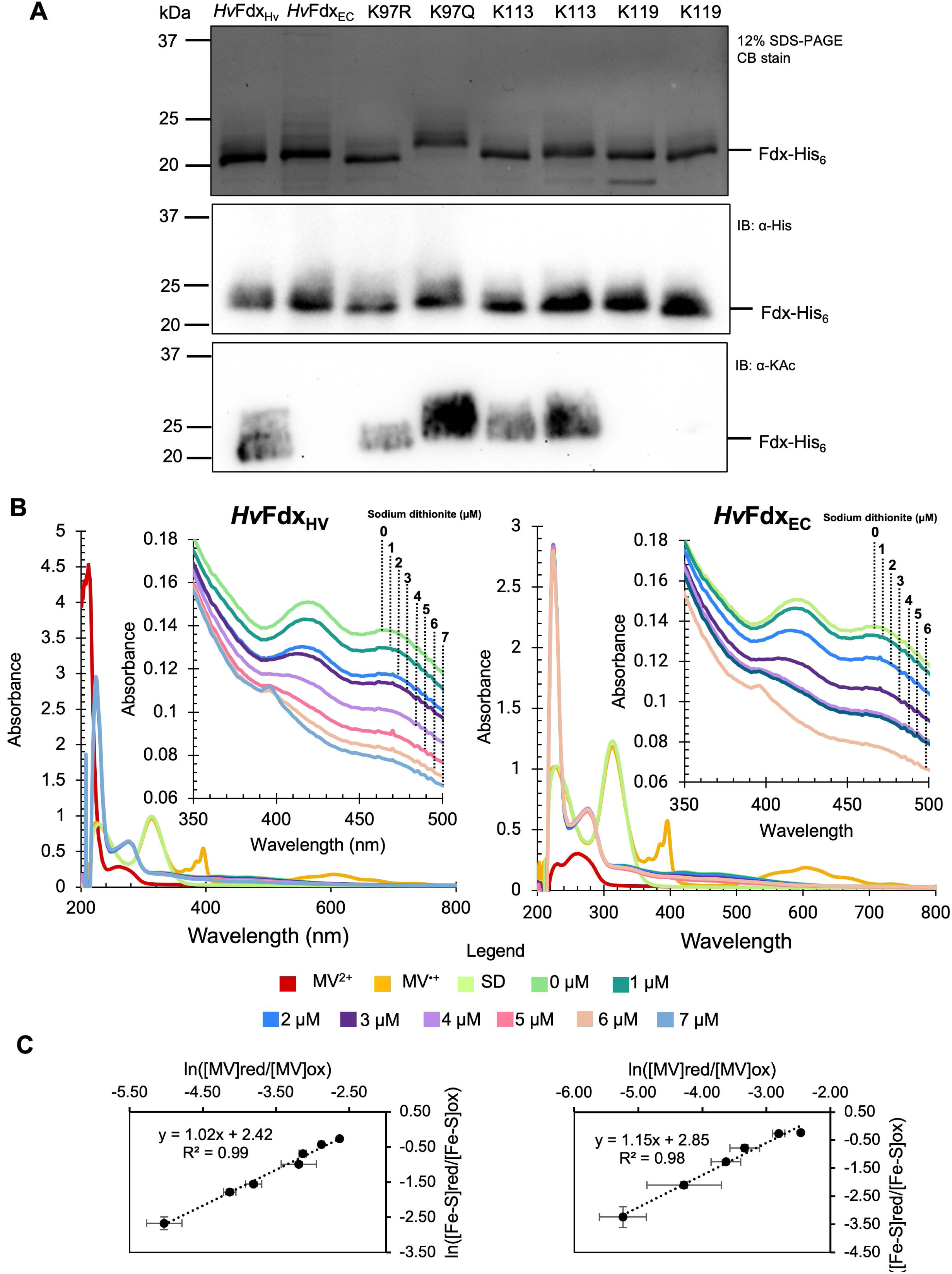
Impact of acetylation on *Hv*Fdx acetylome profile and midpoint. Protein purification was by Ni-NTA affinity and size exclusion chromatography from cells grown in Hv-Ca^+^ medium. Proteins were eluted in 20 mM HEPES [pH 7.5] and 2 M NaCl buffer. A. Proteins were normalized to 1 ug and precipitated with 10% TCA overnight, then boiled for 10 minutes prior to being separated by reducing SDS-PAGE. Wild-type *Hv*Fdx expressed in *H. volcanii* (*Hv*Fdx_HV_) and *E. coli* (*Hv*Fdx_EC_), along with the lysine variants K97R/Q, K113R/Q, and K119R/Q expressed in *H. volcanii*, were purified and separated by reducing 12% SDS-PAGE, and analyzed by Coomassie blue staining (**top**), anti-His tag western blotting (**middle**), and anti-acetyllysine western blotting (**bottom**). SDMs were generated by the reduce recycle PCR (rrPCR) method. WT Fdx exhibited a predicted pI of 4.17. Similarly, K97R, K113R, and K119R variants retained a pI of 4.17, whereas K97Q, K113Q, and K119Q variants displayed a reduced predicted pI of 4.11. B. Oxidized Fdx (20 µM) purified from *H. volcanii* (*Hv*Fdx_HV_) and *E. coli* (*Hv*Fdx_EC_) was reacted anaerobically with increasing concentrations of sodium dithionite (SD) in the presence of 10 µM methyl viologen (MV^2+^) as a redox mediator and reference dye. After 30 min of equilibration, UV–visible spectra were recorded. Values at A_604_ (MV^•^⁺) and A_470_ (Fdx-bound [Fe-S]) were used to calculate the reduction state of the enzyme. C. The midpoint redox potential (E₀′) of Fdx-bound [Fe-S] was determined by fitting the titration data to the modified Nernst equation (see method) by plotting ln([MV_red_]/[MV_ox_]) against ln([Fe-S]_red_]/[Fe-S]_ox_]). Control spectra were collected for 10 µM oxidized MV^2+^, 10 µM fully reduced MV^•+^ (generated with 100 µM SD), and 100 µM SD alone. Each data point represents the mean ± SD of three independent experiments.

### K119 is the primary acetylation site of *Hv*Fdx

To further investigate the acetylation signal observed for the *H. volcanii* purified *Hv*Fdx-His_6_ protein, the lysine residues identified to be acetylated (K97, K113, and K119) by mass spectrometry analysis [6] were targeted for amino acid substitution. Lysine-to-arginine (K→R) substitutions were used to retain the positive charge and mimic the deacetylated state, while lysine-to-glutamine (K→Q) substitutions were used to neutralize charge and mimic the acetylated state. The corresponding *Hv*Fdx-His_6_ variants were expressed in *H. volcanii*, purified, and analyzed in parallel with the wild-type protein (**Fig. 3A**). Relative to wild-type *Hv*Fdx-His_6_, the acetylation signal was strongly reduced or absent in the K119R and K119Q variants, while substitutions at K97 and K113 had little to no effect on lysine acetylation. Thus, K119 is the predominant site of lysine acetylation in *Hv*Fdx under the conditions tested.

### Redox potential of *Hv*Fdx

Following the purification of the [2Fe-2S] coordinated *Hv*Fdx-His_6_ in acetylated and non-acetylated forms, the potential effects of lysine acetylation on the redox properties of *Hv*Fdx were investigated. Iron quantification indicated successful Fe incorporation into *Hv*Fdx-His_6_ variants, as K119R (Fe, 2.19 ± 0.15 µM) and K119Q (Fe, 2.07 ± 0.08 µM) did not display significant differences in iron content compared to WT. Thus, the midpoint redox potential (*E*_0_′) of *Hv*Fdx_HV_-His_6_, representing the acetylated form purified from *H. volcanii*, and *Hv*Fdx_EC_-His_6_, representing the non-acetylated form purified from *E. coli*, was determined by anaerobic reductive titration using sodium dithionite (Na_2_O_4_S_2_, *E*_0_′ = -660 mV, pH 7.0) as the electron donor/titrant and methyl viologen (MV^2+^; *E*_0_′ = -446 mV, pH 7.0) as an electron mediator/reference dye. Following equilibration at each titration point, UV–visible spectra were recorded to monitor the redox states of methyl viologen at A_604_ and *Hv*Fdx at A_470_ (**Fig. 3B**). Incremental addition of sodium dithionite resulted in a progressive decrease in A_470_, consistent with reduction of the *Hv*Fdx-His_6_-bond [Fe–S] cluster. Fitting the experimental data to the rearranged Nernst equation (see method) yielded a midpoint redox potential of -385 ± 11 mV with 1.02 electrons transfer event (*n*) for *Hv*Fdx_HV_-His_6_-bound [Fe-S] and -382 ± 16 mV with 1.15 electrons transfer event (*n*) for *Hv*Fdx_EC_-His_6_-bound [Fe-S] (**Fig. 3C**). These values are consistent with previously reported midpoint potentials for untagged haloarchaeal [2Fe-2S] ferredoxins determined by EPR spectroscopy (−384 mV) where europium(II) chloride (*E*_0_′ = −0.40 V versus the standard hydrogen electrode [SHE]) and europium(II)–EGTA (*E*_0_′ = −0.90 V versus SHE) were used as chemical reductants to reduce the [2Fe–2S] cluster [53]. Collectively, these findings indicate that neither lysine acetylation nor the C-terminal His_6_-tag substantially altered the redox properties of *Hv*Fdx under the conditions tested.

### Impact of lysine acetylation on *Hv*Fdx-mediated electron transfer to DCIP

To investigate the functional consequences of lysine acetylation, the *Hv*Fdx-His_6_ preparations representing distinct acetylation states were analyzed for electron transfer activity using sodium dithionite as the electron donor and 2,6-dichlorophenolindophenol (DCIP, *E*_0_′ = +217 mV) as the electron acceptor (**Fig. 4A**). The electron transfer activities of *Hv*Fdx_HV_-His_6_ purified from *H. volcanii* (acetylated) and *Hv*Fdx_EC_-His_6_ purified from recombinant *E. coli* (non-acetylated) were compared. At low concentrations of *Hv*Fdx_HV_-His_6_ and *Hv*Fdx_EC_-His_6_ (0.3 –1 µM), comparable electron transfer rates (nmoles of DCIP reduced per minute) of the sodium dithionite reduced *Hv*Fdx → DCIP were observed. However, at higher protein concentrations (1.5–3 µM), the electron transfer activity of *Hv*Fdx_EC_-His_6_ approached saturation (*V*_max_), while that of His_6_-*Hv*Fdx_HV_ did not reach saturation and exhibited a 1.7-fold higher rate of activity. Altogether, these findings suggest that lysine acetylation increases the electron transfer activity of *Hv*Fdx to an electron acceptor such as DCIP.

**Figure 4.**
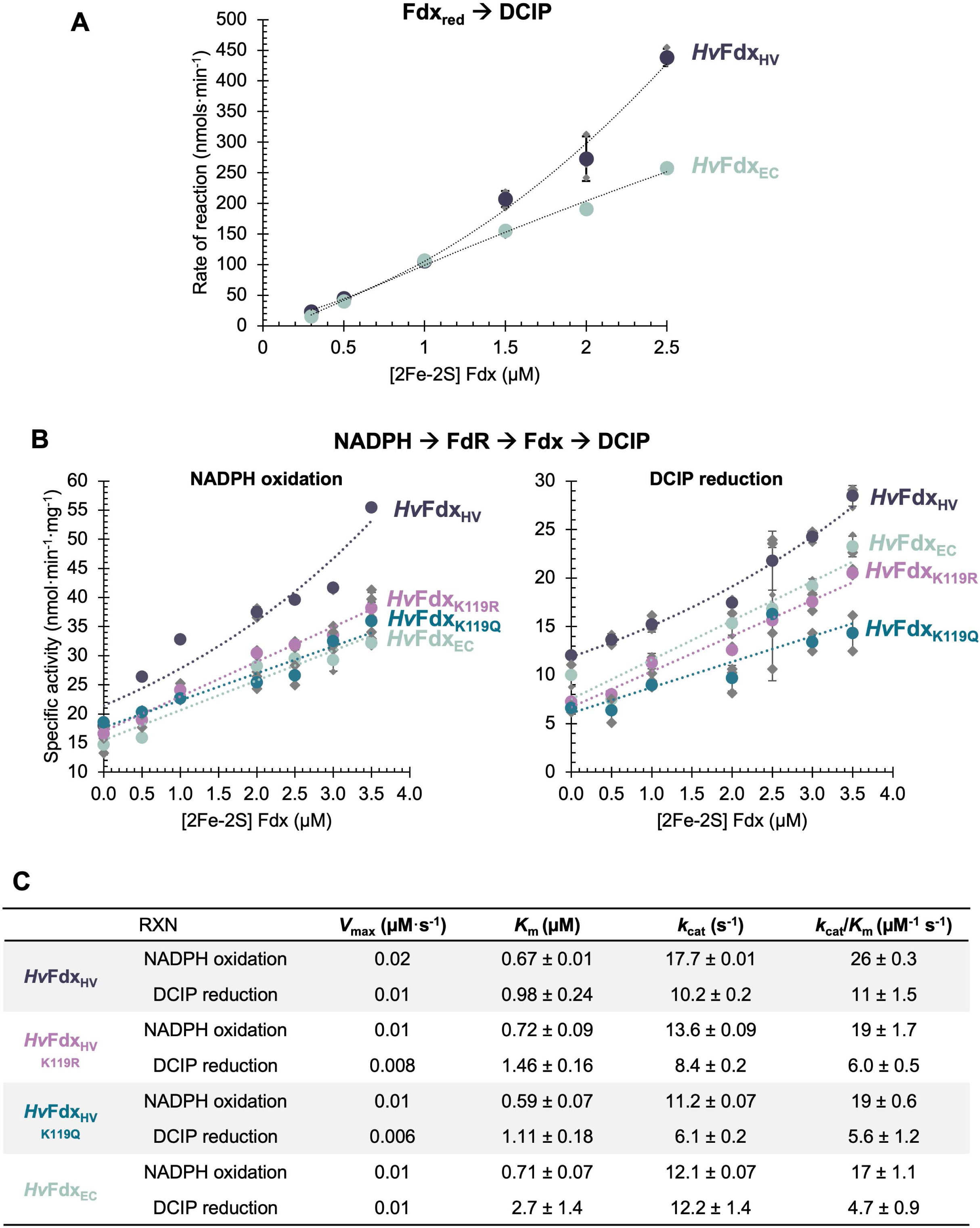
Electron transfer kinetics of *Hv*Fdx under anaerobic conditions. A. DCIP reduction catalyzed *Hv*Fdx. *Hv*Fdx (20 µM) was reduced by sodium dithionite (7 µM). Reactions (100 µL) contained Fdx_red_ (0-2.5 µM), and the artificial electron acceptor DCIP (15 µM) in 20 mM HEPES [pH 7.5] and 2 M NaCl. DCIP reduction was monitored spectrophotometrically at A_600_. Measurements were recorded at 1 min intervals for 30 min at 28 °C. Enzyme rate of reaction was calculated in nanomoles of substrate reduced per minute (nmol·min^-1^). Dashed lines represent the best-fit curves generated from the mean specific activity values. B. NADPH oxidation and DCIP reduction catalyzed by FdR-Fdx. Reactions (100 µL) contained FdR_ox_ (1 nM), Fdx_ox_ (0-3.5 µM), and the artificial electron acceptor DCIP (15 µM) in 20 mM HEPES [pH 7.5] and 2 M NaCl. Reactions were initiated by the addition of NADPH (4 µM). NADPH oxidation and DCIP reduction were monitored spectrophotometrically at A_340_ and A_600_, respectively. Measurements were recorded at 1 min intervals for 30 min at 28 °C. Specific activity was calculated in nanomoles of substrate oxidized or reduced per minute per mg of enzyme (nmol·min^-1^·mg^-1^). At 3.5 μM *Hv*Fdx, the acetylated *Hv*Fdx_HV_ supported significantly greater NADPH oxidation and DCIP reduction than non-acetylated *Hv*Fdx_EC_ (NADPH, *p* = 0.0002; DCIP, *p* = 0.003). Dashed lines represent the best-fit curves generated from the mean specific activity values. C. Steady-state kinetics of NADPH oxidation and DCIP reduction under anerobic conditions. Kinetic parameters include *V*_max_ (µM·s^-1^), *K*_m_ (µM), *k*_cat_ (s^-1^), and *k*_cat_/*K*_m_ (µM^-1^ s^-1^). Values are reported as mean ± standard deviation. Steady-state kinetic parameters were determined from the initial linear portion of each reaction, where less than 20% of the NADPH substrate was consumed.

### Impact of *Hv*Fdx acetylation on linear electron transfer activity

Given the observed effect of lysine acetylation on *Hv*Fdx electron transfer activity, its influence on electron transfer between *Hv*Fdx and its *in vitro* biological partner, *Hv*FdR, was next examined. To monitor electron transfer between these redox partners, assays were performed using purified *Hv*FdR and *Hv*Fdx, with NADPH as the electron donor and DCIP as the electron acceptor. A previous study demonstrates an NADPH → *Hv*FdR → *Hv*Fdx → DCIP electron transfer pathway (Weber *et al*., submitted). *Hv*FdR is required for this activity and addition of *Hv*Fdx stimulates both NADPH oxidation and DCIP reduction in a concentration-dependent manner, confirming functional electron transfer between *Hv*FdR and *Hv*Fdx under the assay conditions tested. Here, the form of *Hv*Fdx acetylated in the native organism (*Hv*Fdx_HV_) is found to support an increase in activity compared to the *Hv*Fdx_K119R_ and *Hv*Fdx_K119Q_ variants and non-acetylated form produced in recombinant *E. coli* (*Hv*Fdx_EC_) (**Fig. 4C**). The naturally acetylated form (*Hv*Fdx_HV_) exhibited the highest *V*_max_ and catalytic efficiency for both NADPH oxidation and DCIP reduction. In contrast, the K119R and K119Q variants displayed reduced *V*_max_, *k*_cat_, and catalytic efficiency in both NADPH oxidation and DCIP reduction, indicating that disruption of K119 acetylated impairs electron transfer. Similarly, the non-acetylated *Hv*Fdx_EC_ exhibited reduced catalytic efficiency compared with *Hv*Fdx_HV_. Although K119R and K119Q are commonly used to mimic the non-acetylated and acetylated states, respectively, these substitutions do not fully replicate lysine acetylation states as arginine contains a larger guanidinium side chain, whereas glutamine possesses a shorter amide side chain than acetylated lysine. Together, these findings indicate that lysine acetylation of *Hv*Fdx enhances electron flux between *Hv*FdR and *Hv*Fdx, promoting more efficient transfer of reducing equivalents from NADPH to the terminal electron acceptor.

### Mapping *Hv*FdR PTM sites onto 3D structural models and their influence on lysine acetylation

Having established that lysine acetylation influences *Hv*Fdx function, the scope of other PTMs within the *Hv*Fdx-*Hv*FdR electron transfer pathway was further assessed by examining the ferredoxin reductase *Hv*FdR (HVO_2345). Specifically, the effects of acetylation at five sites identified by LC-MS/MS analysis (*Hv*FdR K137, K176, K307, K360, and K378) (**Supplemental Dataset S1**) [6] on enzymatic activity, flavin binding and/or thermal stability were investigated. To provide context for this analysis, 3D structural modeling of *Hv*FdR was performed and revealed that several of these acetylation sites are positioned near the predicted ferredoxin- and NADP(H)-binding interface (**Fig. 5A**). These findings suggest that lysine acetylation of *Hv*FdR may regulate its enzymatic activity and/or electron transfer dynamics with putative partners. To further evaluate these predictions, site-directed mutagenesis was performed. Lysine-to-arginine (K→R) and lysine-to-glutamine (K→Q) substitutions were generated at the lysine acetylation sites (K137, K176, K307, K360, and K378). The His_6_-*Hv*FdR wild-type and variant proteins were expressed in *H. volcanii* KT05 (*Δfdr*) and purified by tandem Ni-NTA affinity and size exclusion chromatography. The purified proteins were analyzed by SDS–PAGE, Coomassie staining, and immunoblotting using anti-His and anti-acetyllysine antibodies (**Fig. 5B**). In all variants, the lysine acetylation signal intensity was maintained or even increased relative to wild-type *Hv*FdR, and acetylation remained readily detectable in the quintuple mutant variants (K137R/K176R/K307R/K360R/K378R or K137Q/K176Q/K307Q/K360Q/K378Q). The acetylation signal in the multi-site variants further suggest compensatory acetylation at alternative lysine residues. Consistent with this interpretation, purification of His_6_-*Hv*FdR from recombinant *E. coli* retained detectable acetylation (**Fig. 5B**). Together, these observations suggest that lysine acetylation of *Hv*FdR occurs through non-enzymatic mechanisms that involve extensive, dynamic modification at multiple sites.

**Figure 5.**
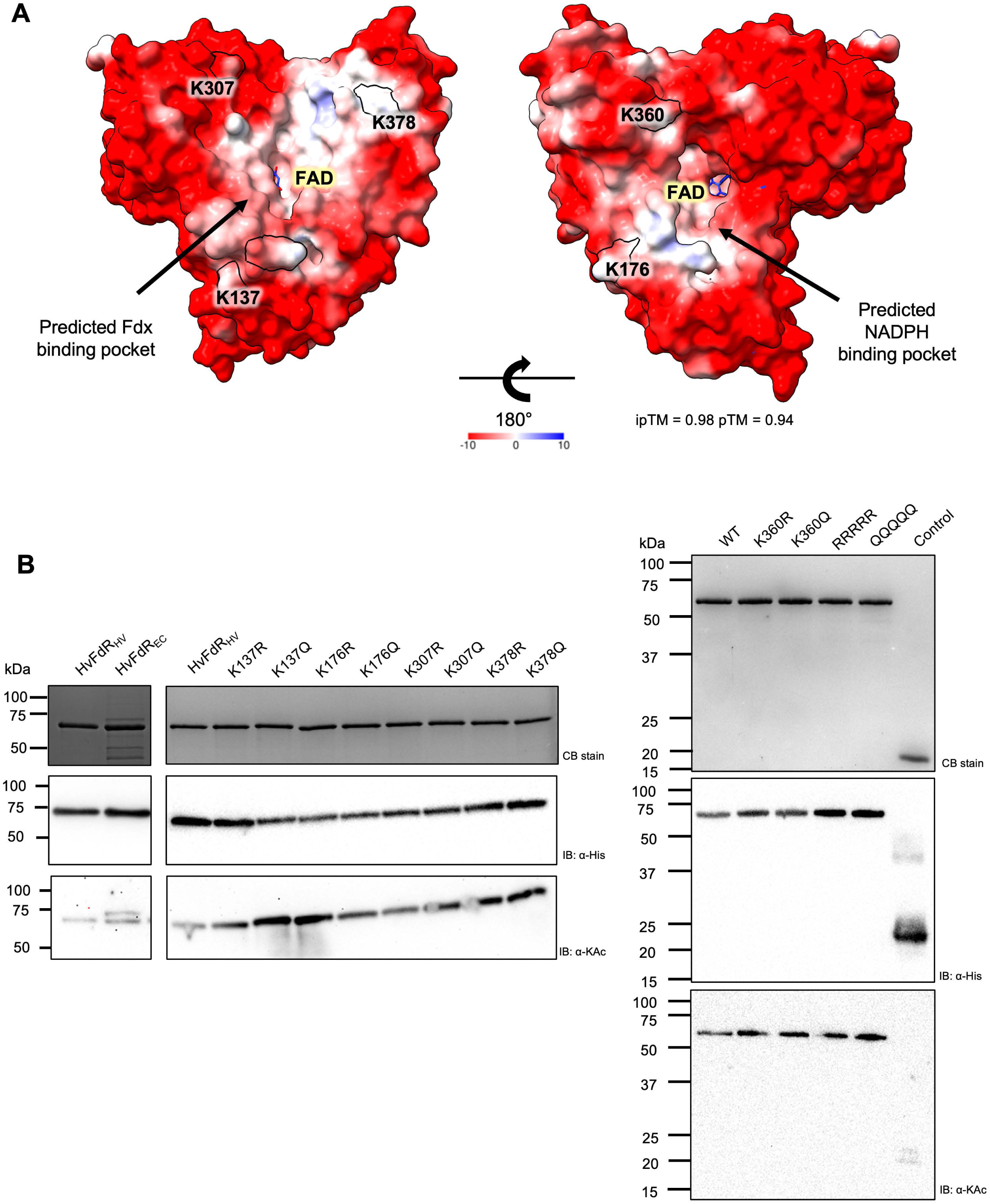
Variants of *Hv*FdR acetylation impact acetylation architecture. A. AlphaFold server predicted model (ipTM = 0.95 pTM = 0.94) with *Hv*FdR from *H. volcanii* with FAD. Electrostatic model of FdR using Alphafold server, with KAc sites K137, K176, K307, K360, and K378. Acetylation at lysine K360 and K378 increases 6.5 × 10^8^ after exposure to hypochlorite of glycerol-grown cells in label free acetylome profile [6]. B. Wild-type *Hv*FdR expressed in *H. volcanii* (*Hv*FdR_HV_) and *E. coli* (*Hv*FdR_EC_) along with the lysine single and quintuple variants expressed in *H. volcanii*, were purified and separated by reducing 12% SDS-PAGE, and analyzed by Coomassie blue staining (top), anti-His tag western blotting (middle), and anti-acetyllysine western blotting (bottom). SDMs were generated by the reduce recycle PCR (rrPCR) method. Control is *Hv*Fdx-His_6_ expressed in H26 Δ*hvo_2874* (KT24), determine to not be acetylated under these conditions.

### Acetylation variants impact on flavin binding of *Hv*FdR

As *Hv*FdR activity depends on incorporation of FAD as a cofactor (Weber *et al*., submitted), the impact of acetylation-site substitutions on flavin binding was evaluated by UV–visible spectroscopy (**Fig. 6A**). Flavin occupancy was estimated by normalizing the characteristic FAD at A_460_ to protein concentration (mg·mL^−1^), and fold changes were calculated relative to wild-type *Hv*FdR. Enzyme concentrations used in subsequent biochemical assays were adjusted according to flavin occupancy (**Supplemental Table S2**). Although all variants exhibited comparable protein concentrations based on A_280_, several acetylation-site variants displayed altered flavin incorporation profiles. The *Hv*FdR K307, K360, K378, and quintuple lysine-substitution variants exhibited increased absorbance at flavin-associated wavelengths (A_380_ and A_460_), suggesting enhanced FAD incorporation or retention relative to wild-type. The K137(R/Q) variants exhibited flavin-binding profiles similar to wild-type, while the K176(R/Q) variants displayed pronounced reductions in flavin occupancy. Despite the observed differences in flavin occupancy, the overall UV–visible spectral profiles were comparable, with no major shifts in flavin-associated absorbance maxima of the variants compared to wild-type. These findings suggest the identified lysine residues influence flavin incorporation or stability without substantially altering the overall flavin-binding environment of *Hv*FdR.

**Figure 6.**
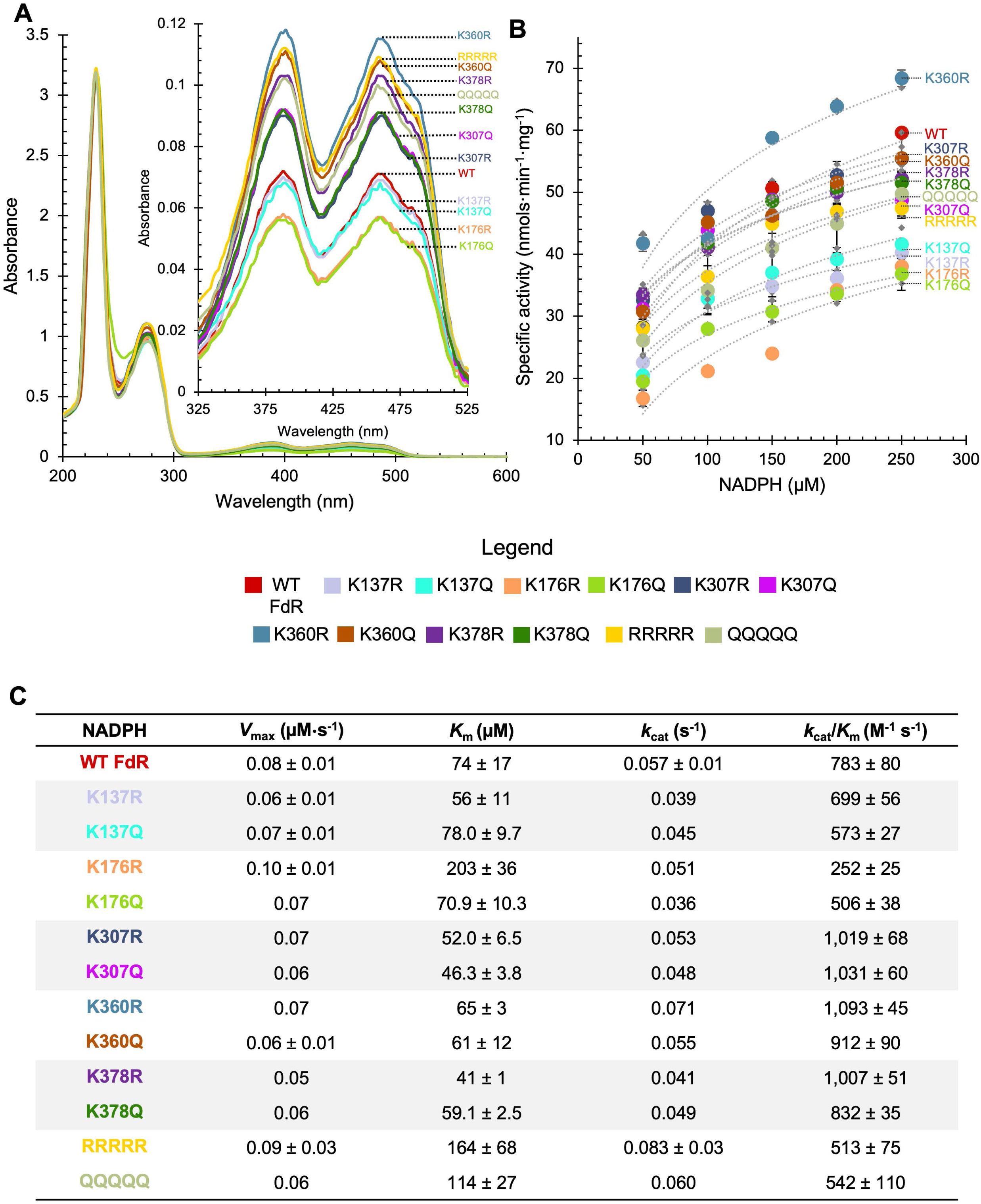
Flavin binding and specific activity of lysine acetylation amino acid substitutions. A. UV-visible spectrum (UVVIS), plotted by wavelength versus absorbance, of His_6_-*Hv*FdR and variant proteins. Proteins were normalized to 0.8 mg·mL^-1^ in 1 mL buffer containing 20 mM HEPES [pH 7.5] and 2 M NaCl. Protein absorbance was monitored from 200 to 600 nm aerobically. All His_6_-*Hv*FdR samples were found to exhibit two peaks at 389 nm and 460 nm, typical of other FAD-dependent proteins. B. Specific activity of His_6_-*Hv*FdR with and without amino acid substitutions. Varying amounts of protein (µM) were added to the reaction dependent on flavin occupancy. NADPH oxidation was tested using concentrations of 50 to 250 µM NADPH in a 150 µL reaction buffer containing 20 mM HEPES [pH 8.0] and 4 M NaCl at 51 °C. A_340_ was monitored over 30 min. Specific activity is defined as nmol NAD(P)H oxidized per min per mg of protein. Dashed lines represent the best-fit curves generated from the mean specific activity values. C. Steady-state kinetics of His_6_-*Hv*FdR and post-translational modification variants utilizing NADPH as the electron donor under aerobic conditions. Kinetic parameters include *V*_max_ (µM·s^-1^), *K*_m_ (µM), *k*_cat_ (s^-1^), and *k*_cat_/*K*_m_ (M^-1^ s^-1^). Values are reported as mean ± standard deviation. Steady-state kinetic parameters were determined from the initial linear portion of each reaction, where less than 20% of the NADPH substrate was consumed. WT FdR, red; K137R, light purple; K137Q, turquoise; K176R, orange; K176Q, bright green; K307R, dark blue; K307Q, magenta; K360R, light blue; K360Q, brown; K378R, purple; K378Q, green; RRRRR, yellow; QQQQQ, sage green. RRRRR, K137R/K176R/K307R/K360R/K378R. QQQQQ, K137Q/K176Q/K307Q/K360Q/K378Q.

### Acetylation-associated residues regulate *Hv*FdR activity

Since *Hv*FdR has an apparent role in redox homeostasis, the impact of the lysine acetylation variants on *Hv*FdR activity was investigated using NADPH oxidase assays. Across all *Hv*FdR variants tested, K137(R/Q) and K176(R/Q) displayed a great reduction in specific activity relative to WT. In contrast, K360R displayed consistently higher specific activity than WT, while K360Q remained comparable to wild-type (**Fig. 6B**). Simultaneous substitution of all five lysine residues resulted in an approximately 1.25-fold (K→R) and 1.19-fold (K→Q) reduction in enzymatic activity at 250 µM NADPH (**Fig. 6B**). Steady-state kinetic analysis revealed that most variants exhibited altered *K*_m_ and *k*_cat_ compared with wild-type FdR, resulting in altered catalytic efficiencies (*k*_cat_/*K*_m_). The K137, K176, and multiple-site variants showed the greatest decrease in catalytic efficiency, reflecting impaired NADPH-dependent catalysis (**Fig. 6C**). These findings suggest that lysine acetylation alters the kinetic properties of *Hv*FdR and that PTM-associated lysine residues contribute to optimal enzyme activity. Together, the results support a role for lysine acetylation in regulating *Hv*FdR-dependent electron transfer and redox metabolism in *H. volcanii*.

### *Hv*FdR variants display differential thermal stability

To characterize the structural effects of the acetylation-site substitutions on *Hv*FdR, thermal stability was assessed by differential scanning fluorimetry (DSF). Protein stability was measured in the presence and absence of the ligand NADP⁺, rather than NADPH, to avoid H_2_O_2_ formation. By this approach, all *Hv*FdR proteins were found to display sigmoidal melting curves characteristic of cooperative thermal denaturation (**Fig. 7**). The *Hv*FdR K137Q variant displayed a smaller unfolding transition than the wild-type, suggesting reduced structural changes during thermal denaturation. In contrast, quintuple arginine and glutamine (K137R/K176R/K307R/K360R/K378R or K137Q/K176Q/K307Q/K360Q/K378Q) variants exhibited decreased thermal stability relative to wild-type. Notably, both the quintuple variants, as well as K360Q, exhibited an additional smaller melt peak (> 60 °C), suggesting the presence of multiple confirmation or cofactor bound states (apo- and holo-protein populations). The altered thermal profiles suggest altered unfolding behavior, reduced flavin release, and/or changes in protein conformational dynamics. In contrast, all other *Hv*FdR variants exhibited melting temperatures comparable to the wild-type protein, indicating that these substitutions did not substantially disrupt global protein folding. Addition of NADP^+^ did not significantly alter protein melting temperatures, suggesting that substrate binding does not markedly influence overall thermal stability under the conditions tested. Collectively, these findings indicate that acetylation-mimicking substitutions, particularly at K137 and in the multi-site variants, influence local structural stability and unfolding cooperativity rather than causing widespread destabilization of *Hv*FdR.

**Figure 7.**
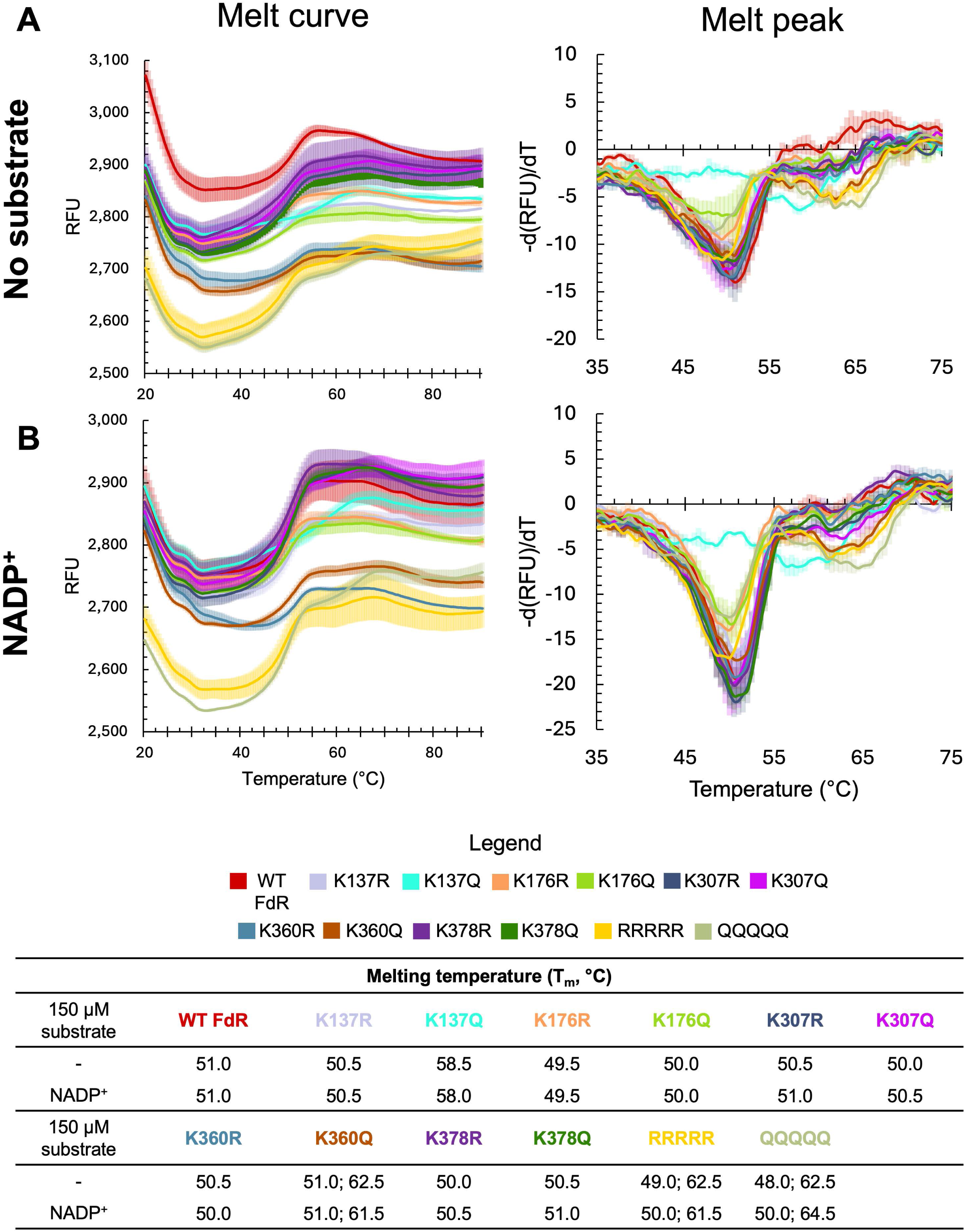
Thermal stability of *Hv*FdR PTM variants. Melting temperatures (T_m_) were determined for WT FdR, as well as lysine single, quadruple, and quintuple variants in buffer containing 20 mM HEPES [pH 8.0] and 4 M NaCl via ThermoFAD methodology. WT FdR, red; K137R, light purple; K137Q, turquoise; K176R, orange; K176Q, bright green; K307R, dark blue; K307Q, magenta; K360R, light blue; K360Q, brown; K378R, purple; K378Q, green; RRRRR, yellow; QQQQQ, sage green. RRRRR, K137R/K176R/K307R/K360R/K378R. QQQQQ, K137Q/K176Q/K307Q/K360Q/K378Q. RFU, relative fluorescence units; d(RFU)/dT, rate at which fluorescence changes as temperature increases. Left, melt curve; right, melt curve. A. Enzyme with no substrate B. Enzyme with 150 µM NADP⁺ C. Melting temperature (°C) determined by peak analysis A student’s *t*-test was used to determine the statistical significance (*p*-value <0.05) of melt peak with WT FdR verse: K137R (0.45), K137Q (4.43 × 10^-4^), K176R (0.13), K176Q (0.15), K307R (0.28), K307Q (0.13), K360R (0.31), K360Q (0.37), K378R (0.18), K378Q (0.18), RRRRR (0.07), and QQQQQ (0.02). A student’s *t*-test was used to determine the statistical significance (*p*-value <0.05) of melt peak with addition of 150 µM NADP^+^ and wild-type FdR verse: K137R (1.0), K137Q (8.43 × 10^-6^), K176R (0.05), K176Q (0.05), K307R (1.0), K307Q (0.05), K360R (0.58), K360Q (0.78), K378R (0.18), K378Q (0.18), RRRRR (0.08), and QQQQQ (0.18). A student’s *t*-test was used to determine the statistical significance (*p*-value <0.05) of melt peak with addition of no substrate added and 150 µM NADP^+^: WT FdR (0.78), K137R (0.39), K137Q (0.05), K176R (0.74), K176Q (0.35), K307R (0.27), K307Q (0.23), K360R (0.23), K360Q (0.38), K378R (0.13), K378Q (0.07), RRRRR (0.65), and QQQQQ (0.02).

### Activity of *Hv*FdR variants with *Hv*Fdx

As acetylation of *Hv*Fdx was observed to impact its electron transfer activity between *Hv*FdR and DCIP, the next objective was to determine whether *Hv*FdR acetylation influences electron transfer between NADPH and *Hv*Fdx. To monitor electron transfer between these redox partners, linear electron transfer assays were performed using purified *Hv*FdR (K137R/K176R/K307R/K360R/K378R or K137Q/K176Q/K307Q/K360Q/K378Q) and the acetylated form of *Hv*Fdx (*Hv*Fdx_HV_), as well as NADPH as the electron donor and DCIP as the electron acceptor. Compared to wild-type *Hv*FdR (**Fig. 4B and 4C**), the quintuple glutamine variant did not impact NADPH oxidation but decreased DCIP reduction by 1.7-fold. The quintuple arginine variant supported a 1.2-fold and 1.1-fold reduction in specific activity of NADPH oxidation and DCIP reduction, respectively (**Fig. 8A**). With kinetic analysis, (**Fig. 8B**), both quintuple arginine and glutamine variants displayed a 1.3- and 1.2-fold reduction in catalytic efficiency, respectively. Together, these findings indicate that lysine acetylation of *Hv*FdR impacts electron flux between NADPH and *Hv*Fdx, promoting efficient transfer of reducing equivalents from NADPH to the terminal electron acceptor.

**Figure 8.**
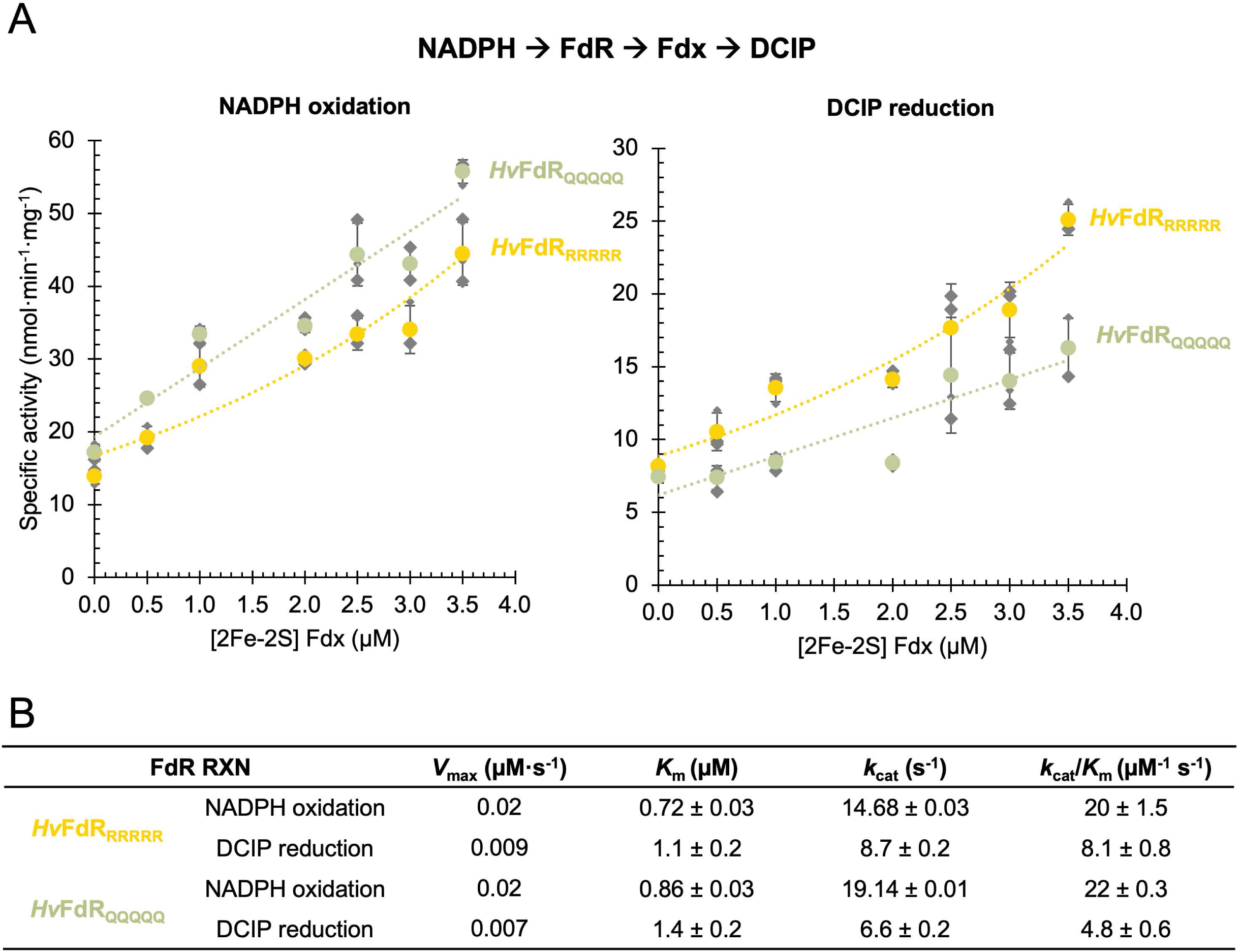
*Hv*FdR acetylation variants impact activity with *Hv*Fdx. A. NADPH oxidation and DCIP reduction catalyzed by FdR-Fdx. Reactions (100 µL) contained FdR_ox_ quintuple variants (1 nM), Fdx_ox_ (0-3.5 µM), and the artificial electron acceptor DCIP (15 µM) in 20 mM HEPES [pH 7.5] and 2 M NaCl. Reactions were initiated by the addition of NADPH (4 µM). NADPH oxidation and DCIP reduction were monitored spectrophotometrically at A_340_ and A_600_, respectively. Measurements were recorded at 1 min intervals for 30 min at 28 °C. Specific activity was calculated in nanomoles of substrate oxidized or reduced per minute per mg of enzyme (nmol·min^-1^·mg^-1^). Dashed lines represent the best-fit curves generated from the mean specific activity values. B. Steady-state kinetics of NADPH oxidation and DCIP reduction under anerobic conditions. Kinetic parameters include *V*_max_ (µM·s^-1^), *K*_m_ (µM), *k*_cat_ (s^-1^), and *k*_cat_/*K*_m_ (µM^-1^ s^-1^). Values are reported as mean ± standard deviation. Steady-state kinetic parameters were determined from the initial linear portion of each reaction, where less than 20% of the NADPH substrate was consumed.

### Identification of HVO_2874 as a GNAT acetyltransferase homolog

Although lysine acetylation of haloarchaeal [2Fe-2S] ferredoxins has been known for more than forty years, the mechanism underlying this modification remains unclear. Here, *Hv*Fdx is lysine acetylated in *H. volcanii* but not in recombinant *E. coli*, suggesting that this modification depends on host-specific enzymatic activity. To identify potential enzymes responsible for lysine acetylation of *Hv*Fdx, 3D-structural modeling was performed between *Hv*Fdx and the 29 candidate GNAT acetyltransferase homologs of *H. volcanii*. HVO_2874 emerged as a strong candidate acetyltransferase, with AlphaFold 3D-modeling predicting its conserved catalytic residue E149 near (4.44 Å) the primary *Hv*Fdx acetylation site, K119 **(Fig. 9A**). GNAT-family acetyltransferases often use conserved glutamate residues as general bases that facilitate deprotonation of the substrate lysine ε-amino group, thereby promoting nucleophilic attack on the acetyl group of acetyl-CoA during catalysis [54–57]. Notably, the primary lysine acetylation site (K119) of *Hv*Fdx is located within a flexible, surface-exposed loop, potentially facilitating accessibility for enzymatic acetylation. Together, these findings suggest that HVO_2874 is a candidate GNAT-family acetyltransferase that may be involved in *Hv*Fdx lysine acetylation.

**Figure 9.**
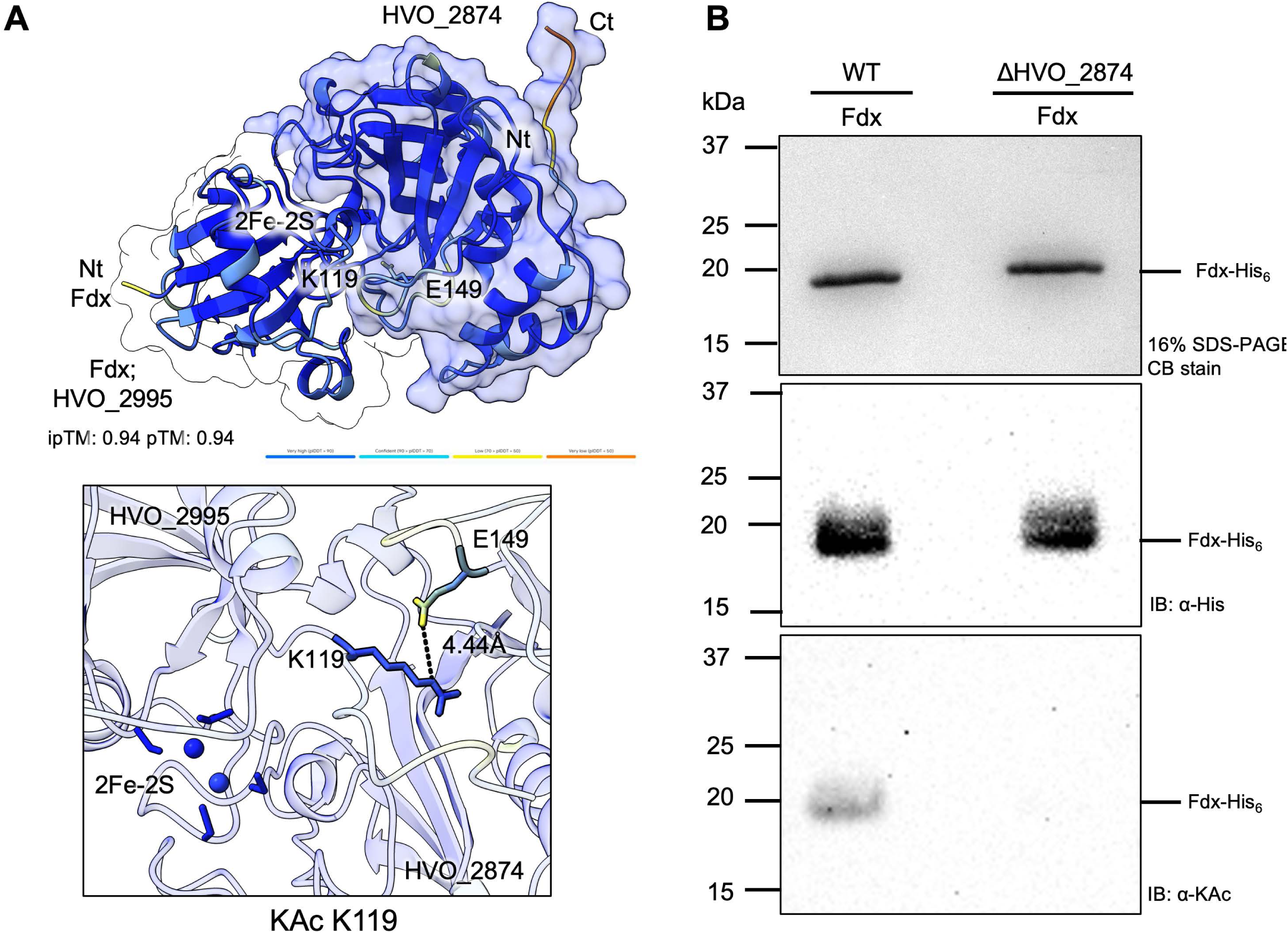
HVO_2874 mediates acetylation of the Fdx (HVO_2995) *in vivo* and *in vitro*. A. AlphaFold-predicted structural interaction between *Hv*Fdx and the putative lysine acetyltransferase *Hv*HVO_2874. The model positions residue K119 of *Hv*Fdx adjacent to the predicted catalytic residue E149 of *Hv*HVO_2874, with an estimated interaction distance of 4.44 Å. *Hv*Fdx is shown containing the 2Fe–2S cluster near K119, and confidence values are indicated by the predicted local distance difference test (pLDDT) color scale. The proximity of K119 to E149 supports the hypothesis that HVO_2874 acetylates *Hv*Fdx at K119. B. *In vivo* analysis of wild-type *Hv*Fdx expressed in wild-type (H26) and H26 Δ*hvo_2874* (KT24) were purified and separated by reducing 16% SDS-PAGE, and analyzed by Coomassie blue staining (top), anti-His tag western blotting (middle), and anti-acetyllysine western blotting (bottom).

### Structural and genomic features of HVO_2874

HVO_2874 contains a disordered N-terminal region (AA 38-63) followed by a predicted N-acetyltransferase domain (AA 67-138, Pfam00583) (**Supplemental Fig. S2A**). Comparative 3D-structural modeling to the characterized GNAT acetyltransferase *Hv*Pat2 [55] identified V147, N125, and Y89 as putative acetyl-CoA binding residues of HVO_2874 (**Supplemental Fig. S2B**). Multiple amino acid sequence alignment and phylogenetic analyses further revealed that HVO_2874 is largely restricted to haloarchaea, a lineage in which 2Fe–2S ferredoxins are extensively acetylated (**Supplemental Fig. S2C**). Across haloarchaeal genera (**Supplemental Fig. S2D-E**), *hvo_2874* homologs were found adjacent to *fen1* (flap endonuclease 1), a DNA replication and repair-associated gene induced under oxidative stress conditions [58, 59]. Conservation of this genomic synteny suggests a functional relationship between lysine acetylation, oxidative stress adaptation, and DNA repair pathways in haloarchaea.

### *hvo_2874* is contributes to lysine acetylation of *Hv*Fdx

Based on structural modeling, the role of HVO_2874 in *Hv*Fdx acetylation was next investigated using a genetic approach. An *H. volcanii* Δ*hvo_2874* mutant strain (KT24) was generated by homologous recombination, and the mutation was confirmed to be specific to the deletion of the *hvo_2874* gene by whole-genome DNA sequencing based on previously reported methods [60]. *Hv*Fdx-His_6_ was then expressed and purified by Ni-NTA affinity and size exclusion chromatography from the wild-type and Δ*hvo_2874* strains. Purified proteins were analyzed by SDS–PAGE, Coomassie staining, and immunoblotting using anti-His and anti-acetyllysine antibodies (**Fig. 9B**). Relative to wild-type, the lysine acetylation signal of *Hv*Fdx-His_6_ was substantially diminished in the Δ*hvo_2874* mutant. Collectively, these findings suggest HVO_2874 contributes to lysine acetylation of *Hv*Fdx either directly or indirectly.

### Global proteomic analysis of the Δ*hvo_2874* mutant reveals loss of lysine acetylation of *Hv*Fdx and other protein targets

To further assess *hvo_2874* for lysine acetylation of *Hv*Fdx and to investigate the broader physiological role of this GNAT acetyltransferase homolog, proteomic analysis was performed. The parental (H26) and Δ*hvo_2874* mutant (KT24) strains were grown to log-phase on minimal medium supplemented with either glycerol or succinate, lactate, and glycerol (SLG) as carbon sources. The proteomes were digested with trypsin, analyzed by label-free quantification (LFQ) LC–MS/MS, and compared for protein identification using FragPipe with MSFragger. Acetylated peptides were identified in the MS-datasets based on a characteristic +42.0106 Da mass shift and classified as either N-terminal or lysine-acetylated peptides according to the modified residue. To ensure high-confidence assignments, acetylated peptides were required to be detected in at least two biological replicates. Proteins displaying complete loss of acetylation intensity in the Δ*hvo_2874* mutant compared to the parental strain were designated as putative HVO_2874 substrates.

Overall, 2,469 proteins were identified in the SLG conditions, whereas 1,335 proteins were detected during growth on glycerol minimal medium (**Supplemental Fig. S3A, Supplemental Dataset S2**). When comparing the parental and Δ*hvo_2874* mutant strains, 233 proteins were differentially expressed under SLG conditions and 11 under glycerol conditions, indicating that loss of *hvo_2874* broadly impacts the proteome. Notably, although *Hv*Fdx (HVO_2995) protein abundance was unaffected by the *Δhvo_2874* mutation, its lysine acetylation, which was detected in the parental strain under both glycerol and SGL conditions, was completely absent in the *Δhvo_2874* mutant. Additional proteins were identified to be lysine acetylated in the wild-type but not the *Δhvo_2874* mutant included glycerol kinase (HVO_1541, *glpK*) and acetyl-CoA synthetase (HVO_A0551, *acs9*) during growth on glycerol, as well as HVO_0769 (TRAM domain protein) and HVO_1797 (beta-lactamase domain-containing protein) during growth on SGL. Among these candidate substrates of HVO_2874, *Hv*Fdx and *Hv*GlpK exhibit the closest predicted interactions with the putative catalytic site active site glutamate residue (E149) of HVO_2874 (**Supplemental Fig. S3B**), consistent with direct enzymatic acetylation. Notably, *Hv*Fdx displayed one of the shortest predicted interaction distances, further supporting *Hv*Fdx as a putative substrate of HVO_2874 (**Supplemental Fig. S3B**).

Interestingly, numerous proteins exhibited N-terminal acetylation in the parental strain that was absent in the *Δhvo_2874* mutant, suggesting that disruption of *hvo_2874* influences PTM systems beyond lysine acetylation. Under SGL conditions, nine proteins were uniquely N-terminally acetylated in the parental strain, whereas 42 proteins exhibited parent-specific N-terminal acetylation during growth on glycerol. Classification according to the acetylated penultimate residue revealed that serine was the predominant target (37 proteins), followed by alanine (10 proteins), glycine (3 proteins), and phenylalanine (1 protein), all of which were acetylated in the parental strain but not in the *Δhvo_2874* mutant (**Supplemental Dataset S2**). Among these proteins, the methionine aminopeptidase homolog HVO_2600 was acetylated on its N-terminal serine residue. This enzyme is predicted to remove the initiator methionine, thereby exposing penultimate residues that serve as substrates for N-terminal acetylation [61]. The nascent polypeptide-associated complex homolog HVO_1382, which is predicted to interact with nascent polypeptides as they emerge from the ribosome [61] was also acetylated on its N-terminal phenylalanine residue. The N-terminal acetylation of both proteins suggests a functional connection between initiator methionine removal and subsequent N-terminal acetylation. Consistent with this model, the ribosomal serine N-acetyltransferase homolog HVO_2425 was more abundant in the parental strain than in the *Δhvo_2874* mutant under SGL conditions (log_2_ fold change = 1.37). The reduced abundance of HVO_2425 in the mutant may contribute to the diminished prevalence of N-terminal serine acetylation observed in this strain.

Overall, our findings demonstrate that HVO_2874 contributes to the lysine acetylation of *Hv*Fdx but does not affect its protein abundance. More broadly, HVO_2874 globally influences both lysine acetylation and N-terminal α-amino acetylation, highlighting its importance in metabolic adaptation and cellular homeostasis.

## Discussion

Post-translational modifications are known to plays roles as central regulators, where lysine acetylation (KAc) functions as a conserved mechanism that links metabolic state to protein activity through fluctuations in intracellular acetyl-CoA and related metabolites [19, 26, 28, 55, 62–65]. Although lysine acetylation has been extensively characterized in bacterial and eukaryotic systems, its role in archaea, specifically archaeal electron transfer systems, remains poorly understood. Here, we demonstrate that lysine acetylation regulates the activity, stability, and electron transfer efficiency of the haloarchaeal ferredoxin (*Hv*Fdx) and ferredoxin reductase (*Hv*FdR), establishing PTMs as modulators of archaeal redox metabolism.

To investigate the functional significance of lysine acetylation in haloarchaeal redox metabolism, the effects of this modification on ferredoxin structure, activity, and electron transfer were examined. Structural modeling positioned *Hv*Fdx K119 proximal to the 2Fe–2S cluster environment and predicted electron transfer interface, suggesting that charge neutralization through acetylation may influence redox partner interactions. Identification of K119 as the predominant *Hv*Fdx acetylation site supports a direct regulatory role for acetylation in ferredoxin function. Redox-active proteins are often influenced by changes in electrostatic environment, structural conformation, and cofactor interactions [33, 36, 66–68]. However, lysine acetylation of *Hv*Fdx did not significantly alter its midpoint potential, Fe–S cluster coordination, or protein abundance, indicating that the thermodynamic driving force for electron transfer and the overall ferredoxin structure remained largely unchanged by this PTM. Instead, acetylation primarily influenced catalytic interactions within the *Hv*FdR–*Hv*Fdx electron transfer pathway, suggesting that the enhanced electron transfer activity arises from changes in the relative orientation rather than changes in the intrinsic redox properties of *Hv*Fdx. Importantly, the measured midpoint potential of the acetylated and non-acetylated forms of *Hv*Fdx-His_6_ closely matched previous EPR-derived values for haloarchaeal ferredoxins (−384 mV), further indicating that the C-terminal His_6_-tag did not substantially impact intrinsic redox properties [53]. Acetylation enhanced electron transfer through the NADPH → *Hv*FdR → *Hv*Fdx → DCIP pathway, supporting a model in which acetylation primarily regulates electron flux rather than fundamentally altering Fe–S cluster chemistry. Similar observations have been reported in the cytochrome P450 CYP101 system, where surface mutations in ferredoxin altered electron transfer efficiency without significantly changing intrinsic cofactor redox properties [69]. Together, these findings suggest that *Hv*Fdx acetylation modulates interaction dynamics between redox partners to optimize electron transfer under hypersaline conditions

While *Hv*Fdx acetylation appears to directly regulate electron transfer interactions, the mechanisms governing *Hv*FdR acetylation remain more complex. Acetylation remained detectable when *Hv*FdR was recombinantly expressed in *E. coli*, suggesting that this *Hv*FdR modification may occur through nonenzymatic mechanisms. Moreover, substitutions of the identified acetylation sites did not diminish overall acetylation levels and, in some cases, resulted in increased acetylation, supporting the presence of compensatory modification sites. Despite this complexity, acetylation-site variants altered *Hv*FdR flavin occupancy, thermal stability, and activity, indicating that acetylation contributes functionally to *Hv*FdR regulation. Several acetylation sites localized near predicted NAD(P)H- and ferredoxin-binding interfaces, suggesting that acetylation may influence electron transfer efficiency by modulating cofactor interactions or protein–protein association dynamics. Thermal stability analyses further revealed that multi-site acetylation variants altered melting temperature profiles (WT, 51 °C; RRRRR, 49 and 62.5 °C; QQQQQ, 48 and 62.5 °C) which suggested unfolding behavior was disrupted and that lysine acetylation has a role in regulating flavoprotein conformational dynamics under hypersaline conditions. Together, these findings suggest that acetylation-associated charge neutralization contributes to fine-tuning *Hv*FdR stability and electron transfer efficiency in high-salt environments where efficient redox coupling is essential for cellular fitness.

The regulatory role of lysine acetylation in *Hv*Fdx and *Hv*FdR is consistent with studies from bacterial and eukaryotic systems demonstrating conserved acetylation-mediated control of redox-active proteins. Acetylation of human cytochrome *c* alters electron transfer efficiency to cytochrome *c* oxidase, while acetylation of *Arabidopsis thaliana* ferredoxin-NADP reductases occurs near the NADP(H)-binding interface and has been proposed to regulate cofactor interactions [33, 36]. Similarly, acetylation sites in *Hv*Fdx and *Hv*FdR localized near redox-active interfaces, supporting the idea that PTM-mediated regulation of electron transfer pathways represents a conserved strategy across domains of life. These findings align with growing evidence that acetylation of redox-active lysine residues modulates protein stability and electron transfer efficiency [70–73].

Beyond its functional impact, genetic and structural analyses identify HVO_2874 as a candidate GNAT-family acetyltransferase associated with *Hv*Fdx acetylation. Expression of *Hv*Fdx in the Δ*hvo_2874* strain displayed a diminished acetylation signal. In addition, recombinant *Hv*Fdx expressed in *E. coli* lacked detectable acetylation, supporting that *Hv*Fdx modification is not primarily driven by spontaneous nonenzymatic acetylation. Proteomic analyses revealed a loss of *Hv*Fdx acetylation in the Δ*hvo_2874* strain, while *Hv*Fdx abundance remained unchanged, supporting a functional rather than abundance-dependent role for acetylation. Beyond *Hv*Fdx, deletion of *hvo_2874* broadly altered the *H. volcanii* acetylome. Proteins associated with diverse cellular processes, including redox homeostasis, central metabolism, transport, DNA replication and repair, stress responses, signal transduction, and PTM systems, increased in abundance supporting a broader role for HVO_2874 in coordinating metabolic and redox-associated protein acetylation.

Interestingly, loss of *hvo_2874* was associated with remodeling of proteins linked to N-terminal acetylation pathways. The ribosomal serine N-acetyltransferase homolog (HVO_2425) was increased in abundance in the parent strain, which may contribute to the observed differences in the N-terminal acetylome. Despite these changes, extensive N-terminal acetylation was retained in both parental and *Δhvo_2874* strains, with the majority of modified proteins shared between strains, suggesting that HVO_2874 primarily influences lysine acetylation of specific proteins (*e.g., Hv*Fdx) rather than serving as a global regulator of N-terminal acetylation. N-terminal acetylation patterns differed substantially between glycerol and SGL growth conditions, supporting a model in which carbon availability shapes acetylation dynamics in haloarchaea. Collectively, these findings indicate that HVO_2874 contributes to broader acetylome remodeling associated with metabolic adaptation, environmental responses, and redox homeostasis.

The relationship between *Hv*Fdx acetylation and carbon metabolism is notable given the central role of ferredoxins in low-potential electron transfer pathways. In *H. volcanii*, acetyl-CoA is generated primarily through pyruvate:ferredoxin oxidoreductase (POR) and AMP-forming acetyl-CoA synthetases (ACSs) [74, 75]. Similar pathways in *Hb. salinarum* utilize ferredoxins for oxidative decarboxylation of 2-oxoacids through pyruvate:ferredoxin oxidoreductase and 2-oxoglutarate:ferredoxin oxidoreductase reactions [76, 77]. Unlike classical NAD^+^-dependent systems, these pathways depend on low-potential ferredoxins to sustain electron transfer. Acetylation of *Hv*Fdx may therefore provide a mechanism for dynamically regulating electron distribution and protein interactions in response to changes in carbon flux or intracellular acetyl-CoA levels. Consistent with a broader regulatory role, lysine acetylation has also been reported for ferredoxins from diverse bacteria and eukaryotes. Among the acetylomes to date [63, 78], a ferredoxin (DR_2075) from *Dinococcus radiodurans* [79] contains an acetylated lysine located proximal to the 2Fe–2S cluster, suggesting that direct regulation of the electron transfer center by acetylation (**Supplemental Fig. S4**)

Collectively, this work establishes *Hv*Fdx and *Hv*FdR as acetylation-regulated redox-active proteins in *H. volcanii* and provides mechanistic insight into how PTMs influence electron transfer pathways in archaea (**Fig. 10**). Our findings demonstrate that PTMs modulate conserved residues associated with electron transfer activity, flavin interactions, and redox-linked function. Identification of HVO_2874 as a candidate acetyltransferase involved in *Hv*Fdx modification supports the possibility of dedicated enzymatic systems regulating redox-active proteins in haloarchaea. Overall, this work also provides insight into how extremophilic microorganisms dynamically regulate metabolism under hypersaline conditions, where altered electron transfer thermodynamics continuously challenge cellular homeostasis. Lysine acetylation may therefore provide a rapid mechanism for fine-tuning electron flow and maintaining redox balance. These findings establish a mechanistic link between protein acetylation and archaeal redox biology and suggest that PTM-mediated regulation of electron transfer pathways may be more widespread across microbial systems.

**Figure 10.**
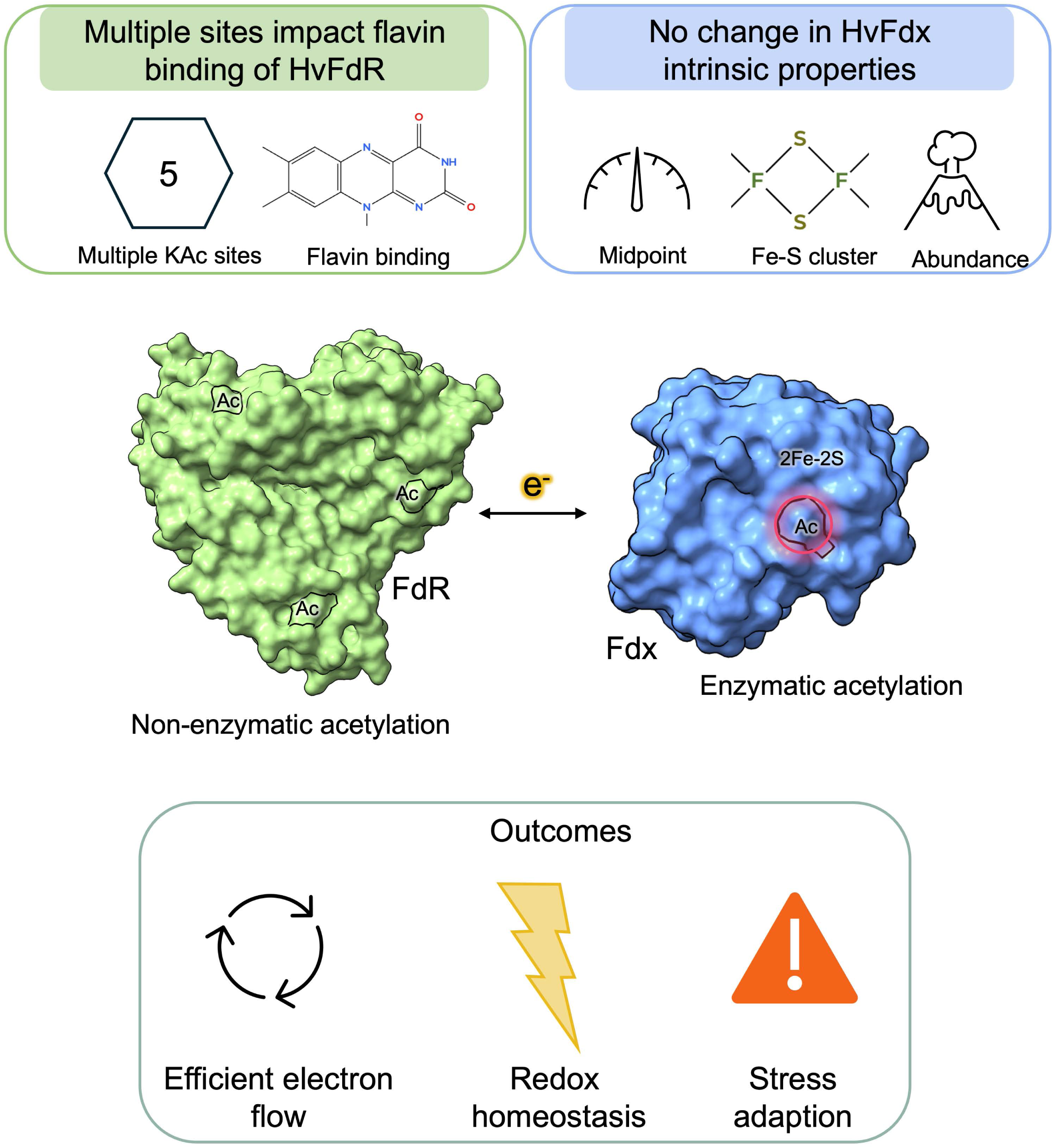
Proposed model for lysine acetylation-mediated regulation of haloarchaeal electron transfer systems. Electrons derived from NADPH are transferred through the FAD cofactor of *Hv*FdR to the 2Fe–2S ferredoxin *Hv*Fdx and subsequently to downstream metabolic or redox-associated electron acceptors. Acetylation of *Hv*Fdx at K119 is associated with altered electron transfer between *Hv*FdR and *Hv*Fdx without substantially altering intrinsic midpoint potential, Fe–S cluster incorporation, or protein abundance of *Hv*Fdx. Potential non-enzymatic acetylation of lysine residues within *Hv*FdR further influences flavin occupancy, enzymatic activity, and thermal stability, suggesting that post-translational modifications regulate protein conformation and electron transfer dynamics. Deletion of the putative GNAT-family acetyltransferase HVO_2874 also resulted in widespread changes to the cellular acetylome and contributes more broadly to lysine acetylation in *H. volcanii*. Overall, lysine acetylation appears to modulate interactions with protein partners to influence electron flow, redox homeostasis, and potentially stress adaptation.

## Materials and Methods

### Materials

Biochemicals and reagents were obtained from Fisher Scientific (Atlanta, GA, USA), Bio-Rad (Hercules, CA, USA), and Sigma-Aldrich (St. Louis, MO, USA). Phusion High-Fidelity DNA Polymerase (cat. no. M0530), restriction enzymes including NdeI (R0111S), BlpI (R0585S), BamHI-HF (R3136S), HindIII-HF (R3104S), and DpnI (R0176), Antarctic phosphatase (M0289S), 2× Quick Ligase (M2200), and KLD enzyme mix (M0554S) were purchased from New England Biolabs (NEB; Ipswich, MA, USA). GeneRuler 1 kb Plus DNA ladder (SM1331) was obtained from Thermo Fisher Scientific (Waltham, MA, USA). Oligonucleotide synthesis and Sanger DNA sequencing were performed by Eurofins Genomics (Louisville, KY, USA).

### Strains and medium conditions

Strains, plasmids, and primers used in this study are listed in **Supplemental Tables S3-4**. *E. coli* strains were cultured at 37 °C in Luria–Bertani (LB) medium supplemented with ampicillin (100 μg·mL^-1^), kanamycin (50 μg·mL^-1^), and/or chloramphenicol (34 μg·mL^-1^). All *H. volcanii* strains were cultivated at 42 °C. ATCC974 medium, pH 6.8, was composed per liter of 125 g NaCl, 50 g MgCl_2_ ·6H_2_O, 5 g K_2_SO_4_, 0.134 g CaCl_2_ ·2H_2_O, 5 g tryptone, and 5 g yeast extract. Minimal medium (Hv-Min) was composed per liter of 144 g NaCl, 18 g MgCl_2_·6H_2_O, 21 g MgSO_4_·7H_2_O, 4.2 g KCl, 441 mg CaCl_2_·2H_2_O, 0.36 mg MnCl_2_·4H_2_O, 0.44 mg ZnSO_4_·7H_2_O, 2.3 mg FeSO_4_·7H_2_O, 0.05 mg CuSO_4_·5H_2_O, 51 mg uracil, 0.8 mg biotin, 0.1 mg thiamine, 10 mM NH_4_Cl or 10 mM urea, 9.75 mL of 0.1 M KPO_4_ buffer [pH 7.5], 0.18% (v/v) glycerol or ‘SLG’ mixture, and 42 mL of 1 M Tris-Cl [pH 7.5] buffer. Carbon sources, 0.18% (v/v) glycerol (GMM) or an ‘SLG’ mixture of 5.5 mM (w/v) succinate acid, 0.1% (v/v) lactic acid, 0.01% (v/v) glycerol, were prepared outlined in the *Halohandbook* [80]. Media were supplemented with novobiocin (0.2 μg·mL^-1^) for plasmid selection or 5-fluoroorotic acid (5-FOA; 50 μg·mL^-1^ dissolved in DMSO) for chromosomal deletion. *H. volcanii* strains were additionally cultured in Hv-Ca^+^ medium supplemented with uracil (50 μg·mL^-1^) according to the Halo Handbook [80]. For solid medium, agar was added to a final concentration of 1.5% (w/v). Strains were stored at −80 °C in 20% (v/v) glycerol stocks. Cryopreserved *H. volcanii* cultures were prepared by diluting stationary-phase cultures 1:1 in 40% glycerol. Cryopreserved *H. volcanii* cultures were prepared by diluting stationary-phase cultures 1:4 in cryopreservation solution consisting of 80 mL glycerol, 20 mL 30% salt water, and 0.2 mL 0.5 M CaCl_2_.

### DNA amplification for plasmid construction

Genomic DNA isolated from *H. volcanii* H26 using the DNA spooling method described in the *Halohandbook* [80] served as the template for PCR amplification of target genes. Amplification reactions were performed using Phusion High-Fidelity DNA Polymerase in GC buffer supplemented with 3% (v/v) DMSO according to the manufacturer’s recommendations (NEB). Reactions were initiated by transferring samples directly from ice to the initial denaturation step (“hot-start” approach) and carried out using a T100 Thermal Cycler (Bio-Rad Laboratories, Hercules, CA, USA). PCR amplicons and DNA restriction fragments were resolved by electrophoresis on 0.8% (w/v) agarose gels. DNA fragments were purified using Monarch PCR & DNA Cleanup or Monarch Gel Extraction kits (NEB, Ipswich, MA, USA). Recombinant plasmids were propagated in *Escherichia coli* Top10 and subsequently GM2163 cells and selected on LB agar supplemented with ampicillin (100 μg·mL^-1^) [81, 82]. Plasmid DNA was purified using the PureLink Quick Plasmid Miniprep Kit (Invitrogen, Waltham, MA, USA). Primer design, restriction enzyme compatibility, cloning orientation, and coding sequence integrity were evaluated in silico using SnapGene (v8.0) and NEB Cloner software prior to plasmid construction. All plasmid constructs generated in this study were verified by Sanger DNA sequencing (Eurofins Genomics, Louisville, KY, USA).

### Generation of *hvo_2874* deletion strain

The *H. volcanii* Δ*hvo_2874* mutant strain KT24 (H26 Δ*hvo_2345*) was generated using a *pyrE2*-based pop-in/pop-out homologous recombination strategy adapted from previously described methods [83]. To construct the deletion plasmid, approximately 500 bp regions upstream and downstream of *hvo_2874* were PCR amplified and cloned into the BamHI and HindIII sites of pTA131, generating the pre-deletion construct pJAM4473. The *hvo_2874* coding region was subsequently removed by inverse PCR while preserving the flanking genomic regions and neighboring open reading frames. Following amplification, template plasmid DNA was digested with DpnI, and the linear PCR product was purified and circularized using the KLD Enzyme Mix (New England Biolabs) to generate the final deletion plasmid, pJAM4474.

For mutant construction, pJAM4474 was transformed into *H. volcanii* H26, and transformants were selected on Hv-Ca^+^ (uracil-deficient) medium to promote chromosomal integration of the plasmid through homologous recombination (pop-in event). Correct plasmid integration was confirmed by colony PCR. To facilitate plasmid excision and allelic exchange (pop-out event), verified transformants were cultured in ATCC974 medium supplemented with 50 μg·mL^-1^ 5-fluoroorotic acid (5-FOA) and incubated at 42 °C with shaking at 200 rpm for 3–4 days. Serial dilutions of these cultures were plated on ATCC974 [pH 6.8] supplemented with 1.5% agar (w/v) containing 5-FOA to select for plasmid loss.

Candidate deletion mutants were screened by PCR, and isolates containing the expected deletion genotype were repeatedly isolated as single colonies on 5-FOA-containing medium to ensure genetic homogeneity. Deletion of *hvo_2874* verified by whole-genome sequencing according to previously reported methods (**Supplemental Dataset S3**) [60].

### Generation of the *H. volcanii* expression strain

The *fdx* and *hvo_2874* genes were amplified from *H. volcanii* genomic DNA by PCR using primers engineered with restriction sites **(Supplemental Table S4)**. The *fdx* gene was isolated by PCR using *H. volcanii* gDNA as template and a primer pair that include NdeI and Xhol sites to generate pJAM4459. The *fdx* gene was PCR amplified from pJAM4459 that includes NdeI and NotI sites and placed into pTA963 to remove the native N-terminus His_6_, generating pJAM4455. The resulting amplicon and expression vector were digested with the corresponding restriction enzymes to enable directional cloning. Prior to ligation, the linearized vector backbone was treated with Antarctic phosphatase to reduce vector self-ligation. Digested DNA fragments were purified using agarose gel extraction or PCR cleanup procedures as appropriate.

The purified insert was ligated into the digested backbone using Quick Ligase, generating plasmid for expression of target genes. The recombinant plasmid was initially transformed in *E. coli* Top10 cells and subsequently GM2163 cells and selected on LB agar supplemented with ampicillin (100 μg·mL^-1^) [81, 82]. For E. coli recombinant expression, plasmids were transformed into E. coli Top10, followed by Rosetta and selected on LB agar supplemented with kanamycin (100 μg·mL^-1^) and chloramphenicol (34 μg·mL^-1^) for maintenance of the pRARE plasmid. Following plasmid isolation and sequence verification, plasmids were transformed into *H. volcanii* H26 or H1207, and transformants were selected on ATCC974 medium supplemented with novobiocin (0.2 μg·mL^-1^) for constitute promoters and Hv-Ca^+^ for inducible promoters.

### SDM construction

Site-directed mutants (SDMs) were generated using the reduce-recycle PCR (rrPCR) method as previously described [84], utilizing the expression plasmid pJAM3944 (His_6_-*Hv*FdR) or pJAM4455 (*Hv*Fdx-His_6_) as the template. The resulting constructs included single and multi-variants. After verification in *E. coli* Top10, the resulting plasmids were transformed into KT05 for His_6_-FdR and H1207 for His_6_-Fdx-His_6_ and stocked in 20% (v/v) glycerol stocks as stated above.

## CRISPRi meditated gene repression of *fdx*

i. **Plasmid generation**. Initial attempts to delete *fdx* (*ferA5,* HVO_2995) via homologous recombination using the *pyrE2*-based pop-in pop-out method [83] were performed in *H. volcanii* strains H26 and H1207, both in the presence and absence of the complementing plasmid pJAM4453 (*fdx*⁺), as well as on varying media and carbon sources. This method was unsuccessful, therefore a conditional expression system was developed by placing *fdx* under the control of the tryptophan-inducible promoter *PtnaA* (HVO_0009) [39] on the genome. However, all deletion attempts were unsuccessful, suggesting *fdx* may be essential under the tested conditions therefore, CRISPRi techniques were performed with the following modifications [41]. Four 18 bp spacer sequences targeting regions surrounding the predicted TATA box and translational start codon of *fdx* (*ferA5,* HVO_2995) were designed adjacent to compatible PAM sites for CRISPR interference (CRISPRi). Spacer sequences were introduced into the pMA-PQ telRNase anti-2 plasmid by inverse PCR using primers containing flanking regions of homology. Following amplification, PCR products were gel purified and treated with DpnI in the presence of 10× rCutSmart buffer at 37 °C for 4 h to remove methylated template DNA. The resulting linear products were subsequently circularized using the KLD enzyme mix prior to transformation into *E. coli* Top10 cells. Transformants were screened by colony PCR, and amplification products were analyzed on 1.5% (w/v) agarose gels and further validated by Sanger DNA sequencing. To generate the final CRISPRi constructs, the SE anti-2 cassette containing the *fdx*-targeting spacer sequences was excised from pMA-PQ telRNase anti-2 using KpnI and BamHI and cloned into the pJAM202c vector. The recipient vector was linearized by inverse PCR with KpnI and BamHI restriction enzymes on the end for ligation, followed by restriction digestion, treated with Antarctic phosphatase to minimize vector recircularization, and gel purified prior to ligation. Inserts and vector backbones were ligated using Quick Ligase and transformed into *E. coli* Top10 cells. Correct insertion of the *fdx* spacer sequences was confirmed by Sanger sequencing. Verified plasmids were propagated in E. coli GM2163 prior to transformation into *H. volcanii* HV30.Transformants were grown on Hv-Ca^+^ supplemented with 0.2 µg·mL^-1^ novobiocin, 0.0755 mg·mL^-1^ tryptophan, 0.05 mg·mL^-1^ uracil, and 1.5% (w/v) agar.
ii. **Cellular growth for growth curves and RNA extraction.** For CRISPRi mediated repression, HV30, HV30/EV, HV30/pJAM4489, HV30/pJAM4490, HV30/pJAM4491, HV30/pJAM4492, were inoculated from 20 % (v/v) glycerol stocks (−80 °C) onto ATCC974 rich medium supplemented with 1.5 % (w/v) agar and 0.2 μg·mL^-1^ nov for plasmid selection. Single colonies were inoculated into 5 mL ATCC974 rich medium in 13 × 100 mm culture tubes and incubated at 42 °C with rotary shaking at 200 rpm. Cultures were sub-cultured to an optical density at 600 nm (OD_600_) of 0.02 in 5 mL of ATCC974 in 13 × 100 mm culture tubes and incubated under the same conditions.
iii. **Growth curve**. Cells were sub-cultured a third time to OD_600_ 0.02 with 1 mL ATCC974, supplemented with novobiocin for strains containing plasmids, in 1.5 mL Eppendorf tubes. In a 96-well CellPro cell culture plate (Alkali Scientific, FL), 150 µL of subculture was aliquoted into three replicate wells. Each strain included three biological and three technical replicates. Using the BioTek Epoch 2 microplate reader and Gen5 software (Agilent, Santa Clara, CA), cell growth was measured as follows: OD_600_ was measured every 15 min for 99 h, with aeration (double orbital continuous shaking), and temperature setpoint 42 °C. No inoculum controls were included to assess potential background signals from the medium alone.
iv. **RNA extraction.** Cultures were sub-cultured a third time to an OD_600_ of 0.02 in 5 mL of the respective medium in 13 × 100 mm culture tubes incubated under the same conditions. Once cells reached an OD_600_ of 0.5-0.6, 1 mL of each culture was harvested by centrifugation at 5,000 × *g* for 10 minutes at room temperature (RT). RNA extraction was performed using the Quick-RNA MiniPrep Plus Kit (cat. no. R1057, Zymo) according to the manufacturer’s protocol. RNA integrity was assessed via electrophoresis on a 0.8% (w/v) agarose gel containing 2x RNA loading dye (cat. no. B0363S, NEB). Samples were diluted to a final concentration of 0.4 ng·µL^-1^ and stored at 80 °C until further use.
v. **Quantitative reverse transcription PCR (qRT-PCR).** Quantitative reverse transcription PCR (qRT-PCR) was performed using the Luna Universal One-Step RT-qPCR Kit (New England Biolabs) on a CFX96 Real-Time PCR Detection System coupled to a C1000 Thermal Cycler (Bio-Rad). Reverse transcription reactions were carried out at 55 °C for 10 min, followed by an initial denaturation step at 95 °C for 1 min. Amplification was then performed over 40 cycles consisting of 95 °C for 10 s and 56 °C for 30 s with fluorescence acquisition during each cycle. A final extension step was performed at 60 °C for 30 s. To verify amplification specificity, melt curve analysis was conducted by increasing the temperature from 60 °C to 95 °C following an initial denaturation step at 95 °C for 10 s. Reactions producing a single melt peak were considered specific for downstream analysis. Relative transcript abundance was normalized to the housekeeping gene *rpL10* (*hvo_0484*), and fold changes in gene expression were calculated using the 2^−ΔΔCt^ method. Primer efficiencies were determined using genomic DNA as template, and only primer pairs exhibiting amplification efficiencies between 95–105% with R^2^ values between 0.90 and 1.10 were used. Statistical analysis of relative transcript abundance was performed using Student’s *t*-test.

### Protein modeling

Three-dimensional structural models of *Hv*Fdx were generated using two independent computational prediction platforms: AlphaFold Server 3 [85] and Phyre2 [86]. The AlphaFold-predicted structures exhibited high confidence, with predicted Template Modeling (pTM) scores of 0.93 for *Hv*Fdx. To assess the impact of cofactor binding on structural predictions, models were generated with various ligand combinations. The *Hv*Fdx model complexed with two Fe^3+^ ions retained a pTM score of 0.93 and an interface predicted Template Modeling (iPTM) score of 0.95. The interaction model of HVO_2874 and Fdx with an acetyl-group at K119 had a score of ipTM = 0.94 and pTM = 0.94. The interaction model of HVO_2874 with an acetyl-group at K27 and Fdx with an acetyl-group at K119 had a score of ipTM = 0.91 and pTM = 0.93. Phyre2 was employed for homology-based structural modeling, producing the *Hv*Fdx model with 99.8% confidence. All structural visualizations and analyses were conducted using ChimeraX (version 1.9). Electrostatic surface properties were assessed using Coulombic surface coloring in ChimeraX [87], where negatively charged regions are depicted in red, positively charged regions in blue, and neutral regions in white.

## Expression and purification of *Hv*Fdx-His_6_, His_6_-*Hv*FdR, HVO_2874-SII and variants

i. ***Escherichia coli* growth conditions.** Expression constructs encoding *H. volcanii* ferredoxin (*Hv*Fdx-His_6_) in pET24b plasmid were transformed into chemically competent *E. coli* Top10 and then subsequently transformed into chemically competent *E. coli* Rosetta (DE3) cells for protein expression. Transformants were plated on LB agar containing kanamycin (Kan; 50 μg·mL^-1^) for pET24b and chloramphenicol (34 μg·mL^-1^) for maintenance of the pRARE plasmid. Notably, *Hv*Fdx contains three rare codons. Rosetta cells containing pET24b-Fdx grown on LB plates were flooded with 12 mL of LB, and 3 mL of LB cell mixture was used to inoculate 500 mL LB medium with respective antibiotics in 2.8 L Fernbach flasks. Cultures were incubated at 37 °C with shaking at 200 rpm until OD_600_ reached 0.6–0.8. Protein expression was induced by cooling cultures to room temperature (24 °C), adding isopropyl β-D-1-thiogalactopyranoside (IPTG) to a final concentration of 0.4 mM, and incubating overnight at 20 °C with shaking at 200 rpm. Cultures were grown to stationary phase (OD_600_ = 1.2–1.6).
ii. ***H. volcanii* constitutive based plasmids growth conditions.** KT05/pJAM3944 and variant strains were inoculated from 20% (vol/vol) glycerol stocks (−80 °C) onto ATCC974 solid medium (1.5% (w/v) agar) supplemented with novobiocin (0.2 μg·mL^-1^ in H_2_O), and incubated for 4-6 days at 42 °C. A 100 mL starter cultures in ATCC974 were grown for 2–3 days under the same conditions. Aliquots (5 mL) were used to inoculate 500 mL of fresh medium in 2.8 L Fernbach flasks.
iii. ***H. volcanii* inducible based plasmids growth conditions.** H1207/pJAM4455, H26/pJAM4455, and KT24/pJAM4455 strains were inoculated from 20% (vol/vol) glycerol stocks (−80 °C) onto Hv-Ca^+^ solid medium (1.5% (w/v) agar) supplemented with 0.18% glycerol. A 100 mL starter cultures in Hv-Ca^+^ were grown for 1–2 days under the same conditions. Aliquots (5 mL) were used to inoculate 500 mL of fresh medium in 2.8 L Fernbach flasks. Protein expression was induced at mid-log phase by the addition of 2 mM L-tryptophan, followed by continued incubation for 14–16 h at 42 °C [88]. Cells were grown in Hv-Ca^+^ supplemented with 0.18% glycerol or ‘SLG’ mix as specified. For cultures expressing *Hv*Fdx-His_6_, 2 mM ammonium iron (III) citrate and 2 mM L-cysteine were added to support iron–sulfur cluster biosynthesis.
iv. **Harvesting**. Cultures were grown to stationary phase and cells were harvested by centrifugation (2,862 × *g*, 50 min) at room temperature (Sorvall Evolution RC centrifuge, Fiberlite F9-4×1000y rotor). Cell pellets were stored at –80 °C until further use.
v. **Cell lysis for His-tagged proteins.** Cell pellets were resuspended at a ratio of 5 mL lysis buffer per 1 g (wet weight) of cells composed of 20 mM HEPES [pH 7.5], 2 M NaCl, 20 mM imidazole, 5 mM 2-mercaptoethanol, 5 mM MgCl_2_, 4 mM CaCl_2_, DNase I (4 μg·mL^-1^), and cOmplete Mini EDTA-free Protease Inhibitor Cocktail (per 10 mL lysis buffer) (cat. no. 1183617000, Roche).
vi. **Cell lysis for StrepII-tagged proteins**. Cell pellets were resuspended in lysis buffer composed of 20 mM HEPES [pH 7.5], 2 M NaCl, 1 mM tris(2-carboxyethyl)phosphine (TCEP, prepared fresh), 1 mM CaCl_2_, 3 mM MgCl_2_, DNase I (10 μg·mL^-1^), and the same protease inhibitor cocktail as above.
vii. **Disruption of cell pellets.** Cells were disrupted by French press (Glen-Mills, NJ, USA) for a total of five passages (1,500–2,000 psi, minimum high ratio of 140). The lysate was clarified by centrifugation at 10,000 × *g* for 50 minutes at 4 °C (Thermo Fisher Scientific Sorvall Legend XTR, Fiberlite F14-6×250 LE). The resulting supernatant was filtered sequentially through 0.45 μm bottle filters (PES membrane; cat. no. 25-232, GenClone). Clarified lysates were used immediately for downstream protein purification.
viii. **Ni-NTA affinity chromatography.** His_6_-tagged proteins were purified using a Bio-Rad BioLogic DuoFlow fast protein liquid chromatography (FPLC) system equipped with a 1 mL HisTrap column (Cytiva). Prior to purification, the chromatography system was primed with filtered nanopure water at 1 mL·min^−1^ for 10 min. The column was washed with five column volumes of nanopure water and equilibrated with 10 column volumes of binding buffer consisting of 20 mM HEPES [pH 7.5], 2 M NaCl, and 20 mM imidazole. Clarified lysates were loaded onto the column at a flow rate of 0.5 mL·min^−1^. Following sample application, the column was washed with five column volumes of binding buffer, and bound proteins were eluted using 20 mM HEPES [pH 7.5], 2 M NaCl, and 500 mM imidazole at 1 mL·min^−1^.
ix. **StrepII protein purification**. Affinity purification of StrepII tagged proteins was performed using 10 mL centrifuge columns (Thermo Fisher Scientific, cat. no. 89898) packed with 1 mL of Strep-Tactin Superflow Plus Resin slurry (Qiagen, cat. no. 1057979), corresponding to a 0.5 mL bed volume. The resin was equilibrated three times sequentially with 10 bed volumes of nuclease-free water (NPH_2_O), followed by 10 bed volumes of binding/wash buffer containing 20 mM HEPES [pH 7.5], 2 M NaCl, and 1 mM tris(2-carboxyethyl)phosphine (TCEP, prepared fresh). Cell lysate was applied to the equilibrated resin and incubated for 1 hour at 4 °C with gentle rocking. The resin was subsequently washed five times with 10 bed volumes of binding/wash buffer with centrifugation at 500 × g for 30 sec per wash cycle. If additional clarified lysate remained, the binding and washing procedure was repeated. StrepII-tagged proteins were eluted twice using 2 bed volumes of elution buffer consisting of 50 mM HEPES [pH 7.5], 2.0 M NaCl, and 5 mM d-desthiobiotin (DTB). Each elution step was performed by incubating the resin at 4 °C with rocking for 30 min, followed by centrifugation at 500 × *g* for 30 sec.
x. **Dialysis**. Eluted protein fractions were dialyzed overnight at 4 °C against 20 mM HEPES [pH 7.5] and 2 M NaCl using SnakeSkin dialysis tubing (3.5 kDa molecular weight cutoff; Thermo Fisher Scientific). Dialysis was performed using approximately 2 L of buffer per 1 mL of protein sample.
xi. **Size exclusion chromatography (SEC).** All proteins were further purified by size exclusion chromatography using the BioLogic DuoFlow chromatography system. Prior to loading, protein samples were filtered through 0.22 µm PES syringe filters (Whatman Uniflo, Cytiva) to remove insoluble material. A Superdex 75 10/300 GL column (GE Healthcare) was equilibrated with two column volumes of filtered nanopure water followed by two column volumes of SEC buffer containing 20 mM HEPES [pH 7.5], 2 M NaCl, and freshly prepared 1 mM dithiothreitol (DTT). Approximately 500 µL of protein sample was loaded onto the column and eluted at a flow rate of 0.3 mL·min^−1^ while monitoring A_280_. Molecular mass estimates were determined using a Gel Filtration Standards kit (Bio-Rad) containing t bovine thyroglobulin (670 kDa), bovine Ɣ-globulin (158 kDa), chicken ovalbumin (44 kDa), horse myoglobin (17 kDa), and vitamin B12 (1.325 kDa). Molecular weight calculations were derived from linear regression analysis (R^2^ > 0.99) of the logarithm of molecular mass versus the gel phase distribution coefficient (*K*_av_), calculated as *K*_av_ = (*V*_R_ − *V*_o_)/(*V*_c_ − *V*_o_), where *V*_R_ represents the retention volume, *V*_o_ the void volume, and *V*_c_ the column volume. Fractions containing purified protein were stored at 4 °C and concentrated using Vivaspin 500 centrifugal concentrators (Sartorius) according to the manufacturer’s recommendations.
xii. **UV-Visible spectroscopy (UVVIS).** Proteins samples in 20 mM HEPES [pH 7.5] and 2 M NaCl were loaded into a quartz cuvette (Quartz Spectrophotometer cell Micro 16.50-Q-10/8.5 mm, Bio-Rad). Spectral scans were recorded from 200-600 nm (BioTek Epoch 2, Agilent) aerobically.
xiii. **Aragon gas exchange.** Purified proteins were stored in sealed glass serum vials. To minimize oxidative damage during storage, the vial headspace was exchanged with argon gas five times. Proteins were subsequently stored at 4 °C until further biochemical analyses.

### Iron content

Under aerobic conditions, 100 µL of freshly purified *Hv*Fdx_Hv_ (pJAM4455) and *Hv*Fdx_EC_ (pJAM4459) containing 51 µg of protein in 20 mM HEPES [pH 7.5], 2 M NaCl, and 1 mM DTT (corresponding to 3.3 µM of protein), was mixed thoroughly with 100 µL of 325 mM acetate buffer [pH 4.5] and 50 µL of 10% (w/v) ascorbic acid (prepared in 325 mM [pH 4.5] acetate buffer). The mixture was diluted with 700 µL of NPH_2_O, mixed gently by pipetting, and incubated at 25 °C for 20 min in a water bath. Following incubation, 50 µL of a 10% (w/v) bathophenanthroline solution (prepared in 325 mM acetate [pH 4.5] buffer) was mixed gently into the protein solution by pipetting. A total of 150 µL of the final reaction mix was transferred into a 96-well plate, in triplicates, and was read at A_535_. Control reactions without bathophenanthroline were included to account for the baseline absorbance contributed by Fdx. A standard curve was generated using ferrous ammonium sulfate (Fe(NH_4_)_2_(SO_4_)_2_) at final concentrations ranging from 0 to 50 µM. Based on its molecular composition, 1 µM of (Fe(NH_4_)_2_(SO_4_)_2_) corresponds to 0.143 µM of Fe^2+^. In this assay, Fe^3+^ is separated from the protein by the acetate buffer and reduced to Fe^2+^ by ascorbic acid [89]. The presence of DTT in the reaction buffer contributes to Fe^3+^ reduction [90]. Bathophenanthroline forms a pink-colored complex upon binding to Fe^2+^ at a 3:1 ratio, enabling detection at A_535_ [91].

## Immunoblot analysis of lysine acetylation in *Hv*Fdx-His_6_ and His_6_-*Hv*FdR

**i. Protein concentration.** Protein concentration was quantified using the Quick Start Bradford 1× Dye Reagent (cat. no. 5000205, Bio-Rad) or BCA Protein Assay Kit (cat. no. 23225, Thermo Fisher Scientific) according to the manufacturer’s instructions. Samples of 5 µL were placed in a 96-well plate in triplicate. A standard curve was prepared by diluting BSA (bovine serum albumin) standards in the assay buffer across a range of concentrations. Protein concentrations were determined by comparing the absorbance of the samples to the standard curve.
**ii. TCA precipitation.** Proteins (1 µg) were diluted to a final volume of 100 µL with nanopure water in 1.5 mL screw-cap microcentrifuge tubes. Ice-cold 80% (w/v) trichloroacetic acid (TCA) was added to a final concentration of 10%, followed by gentle mixing and overnight incubation at 4 °C to precipitate proteins. Samples were subsequently centrifuged at 16,000 × *g* for 10 min at 4 °C, and supernatants were carefully removed without disturbing the protein pellets. Pellets were washed with 1 mL ice-cold acetone and centrifuged again at 16,000 × *g* for 10 min at 4 °C. Following removal of the acetone wash, pellets were air-dried at room temperature for approximately 30 min. Protein pellets were resuspended in 1× Laemmli sample buffer supplemented with 10% (v/v) 2-mercaptoethanol and neutralized with 1 M Tris-HCl [pH 8]. Samples were mixed thoroughly by pipetting, boiled for 10 min, briefly centrifuged, and analyzed by SDS-PAGE.
**iii. SDS-PAGE**. Proteins were analyzed by reducing 12-16% sodium dodecyl sulfate-polyacrylamide gel electrophoresis (SDS-PAGE) run at 200 V for 50 min to 1 hr. Precision Plus Protein Kaleidoscope molecular mass marker (Bio-Rad) was used as the molecular weight standard. SDS-PAGE were Coomassie stained (0.2 g of Coomassie Brilliant Blue G-250 in 40% (v/v) ethanol and 5% (v/v) acetic acid) for 1 hour, then destained overnight in deionized water at room temperature. Gels were imaged using the iBright FL 1000 imaging (Thermo Fisher Scientific Scientific) under the visible protein detection setting.
**iv. Membrane transfer for immunoblotting.** Proteins were transferred from unstained SDS-PAGE gels via wet-transfer to a 0.45 µM polyvinylidene difluoride (PVDF) (Immobilon-FL, Millipore-Sigma cat. no. IPFL00010) membrane at 90 V for 150 min at 4 °C using Bio-Rad mini trans-blot chamber according to manufacturer’s instructions.
**v. Anti-his immunoblotting.** Membrane blocking was performed overnight at 4 °C in 50 mL TBS supplemented with 1% (v/v) Tween 20 (TBST) and 5% (w/v) non-fat dry milk. The primary antibody, His_6_ His-tag from mouse monoclonal ab (cat. no. HRP-66005, Proteintech), was added at a 1:10,000 dilution in 25 mL blocking buffer and incubated for 1 h at room temperature. The membrane was washed 5 times with 30 mL TBST.
**vi. Anti-acetyllysine immunoblotting**. Membrane blocking was performed overnight at 4 °C in 50 mL TBS supplemented with TBST and 5% (w/v) BSA. The primary antibody, Pan anti-acetyllysine polyclonal antibody (cat. no. PTM-105, PTM) was added at a 1:4,000 dilution to 20 mL blocking buffer and incubated for 1 h at room temperature. The membrane was washed 5 times for 5 min in ∼30 mL TBS supplemented with TBST. The secondary antibody, mouse anti-rabbit IgG (whole molecule)-HRP-linked antibody (SC-2357, lot #A0318; Cruz Biotech) was diluted 1:10,000 in 25 mL of blocking buffer and incubated with the membrane for 1 h. The membrane was washed 5 times for 5 min in ∼ 30 mL TBST.
**vii. Membrane imagining**. The membrane was covered in 1 mL of ECL Prime (cat. no. RPN2236, Cytiva) according to the manufacturer’s guidelines. The membrane was imaged using the iBright FL 1000 imaging system (Thermo Fisher Scientific) on the chemi-blot setting.

### Functional impact of *Hv*Fdx acetylation

All experiments were conducted under strictly anaerobic conditions within a Coy anaerobic chamber maintained at 98–99% N_2_ and 1–2% H_2_ at 25–27 °C. Prior to chamber equilibration, buffers and enzyme solutions were degassed and exchanged under 100% argon gas. UV–visible absorbance spectra (200–800 nm) were acquired using quartz cuvettes with a 1 cm path length (Spectrophotometer Cell Micro 16.50-Q-10/8.5 mm) on an Agilent BioTek Epoch 2 spectrophotometer equipped with a cuvette holder, unless otherwise specified.

i. **Midpoint potential.** The value of the midpoint redox potential (*E*_0_′) of *Hv*Fdx-bound [Fe-S] was determined by anaerobic reductive titration using sodium dithionite (SD) as the reductant and methyl viologen (MV^2+^) as a redox reference dye. First a series of reaction mixtures (400 µL) containing purified *Hv*Fdx-His_6_ (20 μM), 10 μM MV^2+^, 20 mM HEPES buffer [pH 7.5] and 2 M NaCl was place in 1.5 mL microfuge tubes. Then to these tubes freshly prepared sodium dithionite (SD) solutions of varying concentrations but fixed volumes were added to achieve a final concentration range of 0 - 7 μM and to reduce the [Fe-S] cofactor and the reference dye at increasing degrees. After the reaction mixtures were incubated for 30 min at 27 °C, the respective UVVIS were collected and processed to obtain the absorbance values at A_470_, which is characteristic of oxidized [Fe-S], and at A_604_ for the MV^•+^ radical; the oxidized form (MV^2+^) shows negligible absorbance at A_604_. The UVVIS of the control solutions containing 10 μM MV^2+^ or MV^•+^, and 100 μM SD were recorded in triplicate to verify the spectral features of each component and to confirm the integrity of the SD dilution. The extinction coefficient of the oxidized form of *Hv*Fdx-His_6_ purified from *E. coli* and *H. volcanii* at A_470_ was determined to be 0.0068 μM^-1^·cm^-1^ and 0.0069 μM^-1^·cm^-1^, respectively; the extinction coefficient of reduced methyl viologen radical at A_604_ is 0.012 μM^-1^·cm^-1^ [92, 93]. Using these extinction coefficient values, the concentrations of oxidized [Fe-S] in Fdx and reduced methyl viologen with various SD concentrations were calculated using the following equations:

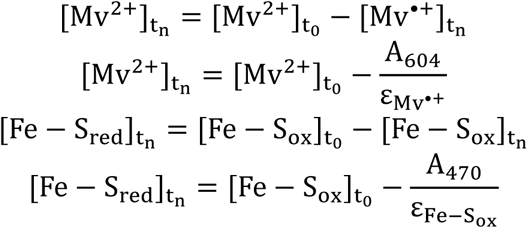

where *t_n_* represents the concentration of the compound at *n* μM SD, and *t_0_* represents the concentration of the compound at 0 μM SD. Then, these values were fitted to the Nernst equation to obtain the midpoint redox potential of *Hv*Fdx-bound [Fe-S], considering that methyl viologen, the redox indicator dye (D), was in equilibrium with the [Fe-S] (Fe-S) [94].

### Nernst equation

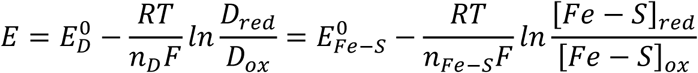

### Rearranged Nernst equation

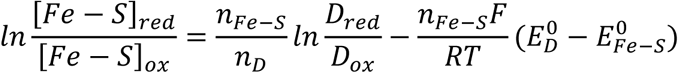

[From the plot of the rearranged Nernst equation: With the value of the slope, 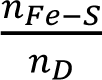, and n_D_ of 1 for MV^2+^/MV^•+^ pair, the electron transfer event for [Fe-S] (*n_Fe-S_*) was determined. The value of the intercept, 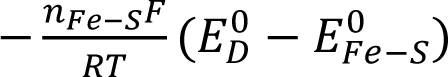, was used to calculate the midpoint redox potential value for Fdx-bound [Fe-S]; *E*_0_′ MV^2+^/MV^•+^, -446 mV vs SHE; gas constant R, 8.314 J K^−1^·mol^−1^; temperature T, 298 K; Faraday’s constant F, 96,485 c·mol^-1^.]

ii. **Molar extinction coefficient of *Hv*Fdx.** The molar extinction coefficient (ε) of the [Fe-S] *Hv*Fdx-His_6_ was determined at A_470_ by plotting absorbance versus protein concentration (0–40 μM) and applying a linear regression to obtain the line of best fit. Measurements were performed in 20 mM HEPES [pH 7.5] containing 2 M NaCl, and monitored by UVVIS. The ε^470^ of the purified *Hv*Fdx-His_6_ was determined to be 5,700 M^−1^·cm^−1^.
  i. ***Hv*Fdx mediated reduction of DCIP**. *Hv*FdR_HV_-His_6_ and *Hv*FdR_EC_-His_6_ (20 μM) was reduced with 7 μM sodium dithionite for 30 min in 20 mM HEPES [pH 7.5] buffer containing 2 M NaCl at room temperature. Excess reductant was removed using a Zeba desalting column following manufacturer’s instructions. To a total reaction (100 μL) in a 96-well plate, Fdx_red_ (0 – 2.5 μM) was added to start the reaction to DCIP (15 μM) were in a reaction buffer composed of 20 mM HEPES [pH 7.5] and 2 M NaCl. DCIP reduction was monitored at A_600_, every minute for 30 min at 28 °C. Rate of reaction (nanomolar of DCIP reduced per minute) and kinetics were determined.
  ii. **Impact of *Hv*Fdx acetylation on electron transfer efficiency to *Hv*FdR.** To a total reaction (100 μL) in a 96-well plate, oxidized *Hv*FdR (1 nM) and *Hv*Fdx (0 – 3.5 μM), were added together and incubated for 2 min at room temperature in a reaction buffer of 20 mM HEPES and 2 M NaCl. The artificial electron acceptor 2,6-dichlorophenolindophenol (DCIP) (15 μM) was added to the protein mix. To start the reaction, NADPH (4 μM) was added. NADP(H) oxidation was monitored at A_340_ and A_600_ for DCIP reduction every minute for 30 minutes at 28 °C.

### Kinetic analysis

Specific activity was calculated as nmol substrate oxidized per minute per mg of enzyme (nmol·min^-1^·mg^-1^), and kinetic parameters were determined by Michaelis–Menten and Lineweaver–Burk analyses. Steady-state kinetic parameters were determined from the initial linear portion of each reaction, where less than 20% of the NADPH substrate was consumed or DCIP was reduced. The Michaelis–Menten curve represents the nonlinear least-squares fit of the experimental data to the Michaelis–Menten equation, 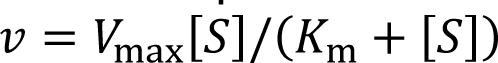 Nonlinear regression was performed using the Solver add-in in Microsoft Excel [95].

## Functional impact of *Hv*FdR acetylation

i. **Protein quantification.** Concentration was determined by measuring UVVIS at A_280_ and calculating the protein extinction coefficient of His_6_-*Hv*FdR. The molar extinction coefficient (ε) was calculated based on the amino acid composition of the protein, specifically the number of tryptophan, tyrosine, and cysteine residues (eq. 1). For concentrations reported in mg·mL^-1^, the molecular weight of the protein was incorporated using where A_280_ is the absorbance at 280 nm, MW is the molecular weight of the protein, and ε_molar_ is the molar extinction coefficient (eq. 2).

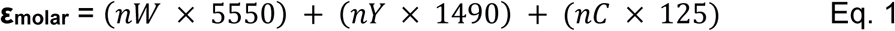

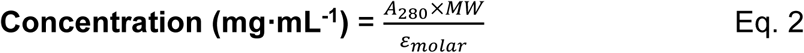

ii. **NAD(P)H oxidase electron transfer assay.** His_6_-*Hv*FdR and acetylation site variants were purified by Ni-NTA affinity chromatography followed by size exclusion chromatography as described above. Purified proteins were analyzed for NADPH oxidase activity under aerobic conditions. Flavin incorporation was estimated by normalizing A_460_ to protein concentration determined from A_280_ measurements, as previously described (Weber *et al*., submitted). Relative flavin occupancy values were calculated for each variant and normalized to wild-type *Hv*FdR to account for differences in flavin loading prior to enzymatic analysis. Prior to enzymatic analysis, proteins were buffer exchanged into 20 mM HEPES [pH 8.0] containing 4 M NaCl using Zeba Spin desalting columns. NADPH oxidase assays were performed using normalized concentrations of *Hv*FdR or variant proteins in the presence of varying concentrations of NADPH (0–250 µM). Reactions were initiated by the addition of NADPH, and NADPH oxidation was monitored spectrophotometrically at A_340_ over 30 min at 51 °C.
iii. **Impact of *Hv*FdR acetylation on electron transfer efficiency to *Hv*Fdx.** To a total reaction (100 μL) in a 96-well plate, oxidized *Hv*FdR (1 nM) varients and wild-type *Hv*Fdx (0 – 3.5 μM), were added together and incubated for 2 min at room temperature in a reaction buffer of 20 mM HEPES and 2 M NaCl. The artificial electron acceptor 2,6-dichlorophenolindophenol (DCIP) (15 μM) was added to the protein mix. To start the reaction, saturating NADPH (4 μM) was added. NADP(H) oxidation was monitored at A_340_ and A_600_ for DCIP reduction every minute for 30 minutes at 28 °C.
iv. **ThermoFAD.** Prior to thermal stability analysis, His_6_-*Hv*FdR and variant proteins were buffer exchanged into 20 mM HEPES [pH 8.0] containing 4 M NaCl using Zeba Spin desalting columns. Protein samples were adjusted to a final concentration of 0.5 mg·mL^−1^ in a total reaction volume of 20 µL. When indicated, samples were supplemented with 150 µM NADP^+^ to evaluate ligand-dependent effects on protein stability. Buffer-only controls were included as negative controls. Samples were loaded into 96-well PCR plates (Bio-Rad) and analyzed using a C1000 thermal cycler equipped with a CFX96 real-time fluorescence detection system (Bio-Rad Laboratories). Thermal denaturation was monitored over a temperature range of 20–90 °C with temperature increments of 0.5 °C every 10 s using the SYBR Green filter setting. Fluorescence intensity data were analyzed to determine the melting temperature (T_m_), corresponding to the unfolding transition peak.

## Acetyltransferase HVO_2874 assays

***i. In vivo* acetylation of *Hv*Fdx.** *Hv*Fdx-His_6_ expression construct was transformed into *H. volcanii* H26 and the H26 Δ*hvo_2874* (KT24) strain. All methods are as described above. Cultures were grown in Hv-Ca^+^ medium supplemented with ‘SLG’ as the carbon source. Cells were harvested by centrifugation and lysed, and *Hv*Fdx was purified using nickel affinity chromatography with a HisTrap column, followed by size exclusion chromatography. Purified protein fractions corresponding to *Hv*Fdx were pooled and concentrated for downstream analyses. For immunoblot analysis, 1 μg of purified *Hv*Fdx from H26 and KT24 were separated by SDS-PAGE under denaturing conditions. Following electrophoresis, proteins were either stained with Coomassie Brilliant Blue or transferred to a PVDF membrane for anti-his and anti-acetyllysine western blot analysis. Antibody incubation, washing, and detection procedures were performed as described above.

## Whole cell acetylome profile of H26 and *hvo_2874* deletion

**i. Cellular growth.** FASP protocol was performed as described [96] with modifications. *H. volcanii* strains (H26 and H26 Δ*hvo_2874*) were inoculated from 20% glycerol stocks (−80 °C) onto ATCC974 rich medium supplemented with 1.5% (w/v) agar and incubated for 5 days at 42 °C. Isolated colonies were transferred to fresh liquid 5 mL Hv-Min supplemented with 0.18% glycerol or ‘SLG’ mixture. Cultures were grown to log phase (OD_600_ of 0.4) at 42 °C in rotating culture tubes at 200 rpm. The cultures were sub-cultured twice to OD_600_ of 0.02 in respective 5 mL Hv-min and grown to log phase at 42 °C in rotating culture tubes. Once the cells were grown to log phase OD_600_ of 0.4, the 5 mL cell culture was harvested by centrifugation (5,000 g x 10 min). The supernatant was removed, and the cell pellet was stored at 80 °C until further use. Three biological replicates were used for an *n* = 3.
**ii. Cell lysis.** Cell pellets were resuspended in 400 μL SDS lysis buffer containing 4% (w/v) SDS in 100 mM Tris-HCl [pH 7.6] and lysed by brief sonication at room temperature to minimize SDS precipitation. Lysates were clarified by centrifugation at 16,000 × *g* for 5 min, and supernatants containing soluble protein were collected. Protein concentrations were determined using the bicinchoninic acid (BCA) assay (Thermo Fisher Scientific) with bovine serum albumin (BSA) as the standard. Samples were reduced by addition of 1 M dithiothreitol (DTT) to a final concentration of 0.1 M.
**iii. Filter-aided sample preparation (FASP) for proteomic analysis.** Up to 50 µg of total protein was diluted in 200 μL UA buffer (8 M urea in 100 mM Tris-HCl [pH 8.5]) and loaded onto 10 kDa molecular weight cutoff filter units (Thermo Fisher Scientific). All centrifugation steps were performed at 24°C to prevent SDS precipitation. Samples were washed with 200 μL UA buffer thrice and alkylated with 100 μL iodoacetamide (IAA) buffer (0.05 M IAA in UA buffer) for 20 min in the dark at room temperature. Filters were subsequently washed trice with 100 μL UA buffer followed by 100 μL ammonium bicarbonate (ABC) buffer thrice to remove residual detergents and alkylating agents.
**iv. Trypsin digestion.** Proteolytic digestion was performed directly on the filter units with ABC buffer (40 μL) and Trypsin Gold (Promega) at an enzyme-to-protein ratio of 1:50 and incubated at 37 °C for 90 min. Resulting peptides were recovered by centrifugation, followed by an additional elution step of 50 μL 0.5 M NaCl. Peptide desalting was followed below.
**v. Peptide desalting and preparation for LC-MS/MS.** Recovered peptides were desalted using C18 spin columns (Thermo Fisher Scientific) with minor modifications to the manufacturer’s protocol. Samples were acidified to 0.2% trifluoroacetic acid (TFA) and adjusted to approximately pH 2 prior to desalting. Columns were equilibrated with 50% acetonitrile (ACN) followed by 0.1% TFA. Peptides were bound to the C18 resin by centrifugation, washed with 0.1% TFA, and eluted using buffer containing 0.1% formic acid (FA) and 80% ACN. Eluted peptides were dried to completion using a SpeedVac concentrator. Dried peptide samples were resuspended in 25 μL 0.1% FA (Buffer A) with gentle agitation and clarified by centrifugation at 16,000 × *g* for 20 min at 4 °C. Supernatants (20 μL) were transferred to LC-MS sample vials, and peptide concentrations were estimated spectrophotometrically by A_280_ using a NanoDrop instrument.
**vi. Mass spectrometry.** Liquid chromatography-tandem mass spectrometry was performed using a Vanquish Neo UHPLC system coupled to an Orbitrap Exploris 240 mass spectrometer (Thermo Fisher Scientific, USA) with a 1-hour LC gradient [97] with the following modifications. 200 ng peptides were loaded onto a 50 cm C18 capillary column and separated using a linear gradient of Solution B (80% acetonitrile, 0.1% formic acid), beginning at 2% and increasing to 4% over 30 sec, followed by sequential increases to 28% over 40 min, 42% over 8 min, and 55% over 4 min. A final wash step at 99% Solution B was performed to ensure complete elution of bound peptides. MS data were acquired in data-dependent mode, with MS1 scans at the resolution of 120k and MS2 fragmentation at 15k resolution. From cells grown in glycerol minimal medium, the precursor ion corresponding to the *Hv*Fdx peptide IVYNAKHLDYLQNR (m/z 596.9846) was selectively isolated from the MS1 scan and subjected to MS2 fragmentation for peptide identification and confirmation of the acetylation site.
**vii. Data analysis.** Raw Thermo MS/MS datasets were analyzed using the FragPipe proteomics platform (v23.0) with MSFragger (v4.3) for peptide identification and label-free quantification (LFQ). Searches were performed using the LFQ-MBR workflow with default label-free settings unless otherwise specified. Sequence databases included target proteins together with decoy and contaminant entries [10.5281/zenodo.15188230]. Search parameters were configured with a precursor mass tolerance of ±10 ppm. Variable modifications included lysine acetylation (+42.0106 Da; up to two sites per peptide), methionine oxidation (+15.99492 Da), cysteine carbamidomethylation (+57.02146 Da), N-terminal acetylation (+42.01057 Da), lysine sampylation (+114.0429 Da), and lysine succinylation (+100.0160 Da). Quantitative comparisons between experimental groups were performed using MaxLFQ intensity values. FragPipe-analyst was used to determine protein abundance with the following analysis outline: Data type, LFQ; Intensity type, Max LFQ intensity; DE adjusted *p*-value cutoff, 0.05; DE log_2_ fold change cutoff; 0.58; normalization type, medium centered.

## Statistical analysis

All immunoblots were performed at least twice with two independent biological and experimental repeats. Mass spectrometry data were acquired using four biological replicates for samples grown in glycerol and SLG minimal media. Student *t*-test analyses were carried out in Microsoft Excel.

## Data availability

The MS-based proteomic datasets generated in this study are publicly available in the UCSD MassIVE repository (Mass Spectrometry Interactive Virtual Environment, https://massive.ucsd.edu) under the following accession number MSV000102701, MSV000102702, and MSV000102723, and PRIDE (PXD082020). Each dataset includes the raw MS files used for peptide identification and quantification. Sequencing data generated by this project were submitted to the National Center for Biotechnology Information (NCBI) Sequence Read Archive (SRA) and can be found under BioProject accession PRJNA1506097.

## Acknowledgments

Funds awarded to JMF to advance archaeal biocatalysts were through the U.S. Department of Energy, Office of Basic Energy Sciences, Division of Chemical Sciences, Geosciences and Biosciences, Physical Biosciences Program (DE-FG02-05ER15650) and to advance cellular mechanisms were through the National Institutes of Health (R35 MIRA 1R35GM161171-01). USDA National Institute of Food and Agriculture (Hatch Project FLA-MCS-006312) funds to JMF also supported this project. Funds awarded to KRW through NASA Florida Space Grant (FSGC 80NSSC20M0093) to determine functional significance of post-translation modifications in haloarchaea. X.W. was supported by the National Science Foundation grant number 2414925 (Original award number 2042182). We gratefully acknowledge Anita Marchfelder and her laboratory for generously providing the CRISPRi strains and plasmids used in this study.

