## Supplemental Table S1 for "Lysine acetylation-mediated regulation of ferredoxin and ferredoxin reductase redox-active proteins in *Haloferax volcanii*"

**Supplemental Table S1.** Comparison of 3D-structures of *Hv*Fdx models predicted by Phyre2 and AlphaFold to each other and to CryoEM and X-ray crystallographic structures of other related haloarchaeal, bacterial and plant Fdx homologs.

| **Organism** | **Protein** | **PDBe; UniProt ID** | **Pfam** | **AA** | **Seq. Id.** | **Prob.** | **E-value** | **RMSD (atom pairs) – Phyre2[a]** | **RMSD (atom pairs) – Alpha Fold[b]** |
|---|---|---|---|---|---|---|---|---|---|
| *Haloferax volcanii* | FerA5, *Hv*Fdx | - D4GY89 | PF00111 | 129 | - | - | - | 1.176 Å (80); 3.061 Å (128) | |
| *Halobacterium salinarum* | Fer2 | 1e10; P00216 | PF00111 | 129 | 88.2 | 1.00 | 1.11e-17 | 0.860 Å (104); 2.707 Å (128) | 1.115 Å (83); 3.197Å (128) |
| *Haloarcula marismortui* | Fer1 | 1doi; P00217 | PF00111 | 129 | 83.5 | 1.00 | 7.87e-23 | 1.155 Å (80); 3.070 Å (128) | 0.628 Å (125); 0.789 Å (128) |
| *Thermosynechococcus vestitus* | PetF1 | 5aui; P0A3C9 | PF00111 | 98 | 33.8 | 1.00 | 7.29e-9 | 1.142 Å (51); 5.069 Å (90) | 0.711 Å (76); 4.087Å (90) |
| *Anabaena* pcc7119 | PetF | 1qt9; P0A3C8 | PF00111 | 99 | 34.7 | 1.00 | 3.48e-9 | 1.149 Å (45); 6.251 Å (91) | 0.711 Å (76); 4.087 Å (90) |
| *Arabidopsis thaliana* | Fd2 | 4zho; P16972 | PF00111 | 148 | 28.4 | 1.00 | 1.14e-8 | 1.178 Å (51); 5.075 Å (88) | 0.672 Å (72); 4.147 Å (88) |

*Hv*Fdx (2Fe-2S) Phyre2[a] and Alpha Fold[b] generated models compared to each other and to other crystalized homologs as indicated. Root Mean Square Deviation (RMSD) values based on pruned and total atom pairs calculated using ChimeraX. Sequence identity (Seq. Id.), probability (Prob.), and E-value metrics determined by comparison of *Hv*Fdx AlphaFold model to Fdx homologs using Foldseek. -, not applicable; PDBe, PDB entry; UniProt ID, accession number; AA, amino acid length. Phyre2 (confidence in the model: 99.8%). PF00111, 2Fe-2S iron-sulfur cluster binding domain.
