## Supplemental Table S2 for "Lysine acetylation-mediated regulation of ferredoxin and ferredoxin reductase redox-active proteins in *Haloferax volcanii*"

**Supplemental Table S2.** UV-visible spectra of $His_6$-HvFdR and variant proteins.

| $His_6$-*Hv*FdR | A280 | mg/mL | Normalization of $A_{460}$/(mg/mL) | Fold-change WT/variant | μM for reaction |
|---|---|---|---|---|---|
| WT FdR | 0.948 | 0.767 | 0.071 / 0.767 = 0.092 | - | 1.50 |
| K137R | 0.981 | 0.794 | 0.069 / 0.794 = 0.086 | 1.07 | 1.60 |
| K137Q | 0.948 | 0.767 | 0.068 / 0.767 = 0.088 | 1.04 | 1.56 |
| K176R | 0.981 | 0.794 | 0.057 / 0.794 = 0.071 | 1.29 | 1.93 |
| K176Q | 0.983 | 0.796 | 0.057 / 0.796 = 0.071 | 1.29 | 1.93 |
| K307R | 1.00 | 0.810 | 0.090 / 0.810 = 0.111 | 0.83 | 1.24 |
| K307Q | 1.014 | 0.821 | 0.091 / 0.821 = 0.110 | 0.83 | 1.24 |
| K360R | 1.074 | 0.869 | 0.115 / 0.869 = 0.132 | 0.69 | 1.03 |
| K360Q | 1.045 | 0.846 | 0.108 / 0.846 = 0.127 | 0.72 | 1.08 |
| K378R | 1.001 | 0.810 | 0.103 / 0.810 = 0.127 | 0.72 | 1.08 |
| K378Q | 1.008 | 0.816 | 0.091 / 0.816 = 0.111 | 0.83 | 1.24 |
| RRRRR | 1.083 | 0.877 | 0.109 / 0.877 = 0.124 | 0.74 | 1.11 |
| QQQQQ | 0.935 | 0.757 | 0.099 / 0.757 = 0.130 | 0.70 | 1.05 |

Absorbance spectra were collected to determine relative flavin binding and fold changes in flavin occupancy for each $His_6$-HvFdR protein. Quantitative analysis of spectral features was used to calculate the micromolar amount of flavin required for subsequent enzymatic assays.
