## Supplemental Table S3 for "Lysine acetylation-mediated regulation of ferredoxin and ferredoxin reductase redox-active proteins in *Haloferax volcanii*"

### Supplemental Table S3. List of strains and plasmids used in this study.

| Name | Description | Ref. |
|---|---|---|
| **Strains** | | |
| ***Escherichia coli*** | | |
| Top10 | $F^-$ *mcrA* Δ*(mrr-hsdRMS-mcrBC) Φ80lacZΔM*15 Δ*lacX74 recA1 araD139* Δ*(ara leu)* 7697 *galU galK rpsL* ($Str^r$) *endA1 nupG λ-* | Invitrogen |
| GM2163 | $F^-$ *ara-14 leuB6 fhuA31 lacY1 tsx78 glnV44 galK2 galT22 mcrA dcm-6 hisG4 rfbD1 rpsL136 dam13::*Tn*9 xylA5 mtl-1 thi-1 mcrB1 hsdR2* | New England Biolabs |
| Rosetta | $F^-$ *ompT hsdS* B(*rB*$^-$*mB*$^-$) *gal dcm* (DE3) p*RARE* ($Cm^R$) | Novagen |
| ***Haloferax volcanii*** | | |
| DS70 | wild-type isolate DS2 cured of plasmid pHV2 | (95) |
| H26 | DS70 Δ*pyrE2* | (79) |
| H1207 | H26 Δ*pyrE2 pitA*$_{Nph}$ Δ*mrr* | (85) |
| KT05 | H1207 Δ*fdr* | (Weber *et al.*, submitted) |
| KT24 | H26 Δ*hvo_2874* | This study |
| HV30 | Δ*pHV2* Δ*pyrE2* Δ*trpA* Δ*leuB* Δ*cas6b* Δ*cas3* Δ*bgaH* | (38) |
| **Plasmids** | | |
| ***Escherichia coli*** | | |
| pET24b | $Kan^r$; pBR322-derived plasmid containing C-terminus $P_{T7}$-6xHis tag | Novagen |
| ***Haloferax volcanii*** | | |
| pTA131 | $Amp^r$; pBluescript II containing $P_{fdx}$-*pyrE2* | (79) |
| pJAM503 | $Amp^r$; $Nov^r$; *H. volcanii*-*E. coli* shuttle vector with coding sequence for N-terminal $His_6$ tag (fused to N-terminus of HVO_0850) | (96) |
| pJAM809 | $Amp^r$; $Nov^r$; pJAM202c containing $P2_{rrn}$ HVO_1862-StrepII (*KpnI* site inserted upstream of StrepII coding sequence) | (97) |
| pJAM202 | $Amp^r$; $Nov^r$; *HVO-E. coli* shuttle expression plasmid with psmB-$His_6$; β subunit of 20S proteasomes expressed with C-terminal $His_6$ tag | (98) |
| pTA963 | $Amp^r$; $P_{tnaA}$ with N-terminal $His_6$-tag *pyrE2*$^+$*hdrB*$^+$ | (85) |
| pJAM202c | $Amp^r$; $Nov^r$; *H. volcanii*-*E. coli* shuttle $P2_{rrn}$ empty vector | (96) |
| pMA-RQ SEanti2 | $Amp^r$; *E. coli* shuttle vector carrying synthetic promoter p.Syn–5′ tRNA-like element–5′ repeat handle–spacer–3′ tRNA-like element–terminator T7 | (39) |
| pJAM4484 | $Amp^r$; $Nov^r$; pJAM202c carrying tele-icrRNA construct from pMARQ SEanti2 | This study |
| **HVO_2345 (*Hv*FdR)-derived** | | |
| pJAM3944 | $Amp^r$; $Nov^r$; *H. volcanii*-*E. coli* shuttle $P2_{rrn}$ $His_6$-*Hv*FdR expression | (Weber *et al.*, submitted) |
| pJAM4465 | $Amp^r$; pTA131-based *fdr* pre-deletion plasmid | (Weber *et al.*, submitted) |
| pJAM4467 | $Amp^r$; pTA131-based Δf*dr* deletion plasmid | (Weber *et al.*, submitted) |
| pJAM4535 | $Amp^r$; $Nov^r$; *H. volcanii*-*E. coli* shuttle $P2_{rrn}$ $His_6$-*Hv*FdR K137R expression | This study |
| pJAM4536 | $Amp^r$; $Nov^r$; *H. volcanii*-*E. coli* shuttle $P2_{rrn}$ $His_6$-*Hv*FdR K137Q expression | This study |
| pJAM4537 | $Amp^r$; $Nov^r$; *H. volcanii*-*E. coli* shuttle $P2_{rrn}$ $His_6$-*Hv*FdR K176R expression | This study |
| pJAM4538 | $Amp^r$; $Nov^r$; *H. volcanii*-*E. coli* shuttle $P2_{rrn}$ $His_6$-*Hv*FdR K176Q expression | This study |
| pJAM4539 | $Amp^r$; $Nov^r$; *H. volcanii*-*E. coli* shuttle $P2_{rrn}$ $His_6$-*Hv*FdR K307R expression | This study |
| pJAM4540 | $Amp^r$; $Nov^r$; *H. volcanii*-*E. coli* shuttle $P2_{rrn}$ $His_6$-*Hv*FdR K307Q expression | This study |
| pJAM4541 | $Amp^r$; $Nov^r$; *H. volcanii*-*E. coli* shuttle $P2_{rrn}$ $His_6$-*Hv*FdR K360R expression | This study |
| pJAM4542 | $Amp^r$; $Nov^r$; *H. volcanii*-*E. coli* shuttle $P2_{rrn}$ $His_6$-*Hv*FdR K360Q expression | This study |
| pJAM4543 | $Amp^r$; $Nov^r$; *H. volcanii*-*E. coli* shuttle $P2_{rrn}$ $His_6$-*Hv*FdR K378R expression | This study |

| | | |
|---|---|---|
| pJAM4544 | Amp$^r$; Nov$^r$; *H. volcanii*-*E. coli* shuttle P2$_{rrn}$ His$_6$-*Hv*FdR K378Q expression | This study |
| pJAM4545 | Amp$^r$; Nov$^r$; *H. volcanii*-*E. coli* shuttle P2$_{rrn}$ His$_6$-*Hv*FdR K137R/K176R/K307R/K360R/K378R expression | This study |
| pJAM4546 | Amp$^r$; Nov$^r$; *H. volcanii*-*E. coli* shuttle P2$_{rrn}$ His$_6$-*Hv*FdR K137Q/K176Q/K307Q/K360Q/K378Q expression | This study |
| **HVO_2995 (*Hv*Fdx)-derived** | | |
| pJAM4455 | Amp$^r$; *H. volcanii*-*E. coli* shuttle P$_{tnaA}$ *Hv*Fdx-His$_6$ expression | This study |
| pJAM4459 | Kan$^r$; *E. coli* shuttle P$_{T7}$ shuttle *Hv*Fdx-His$_6$ expression | This study |
| pJAM4485 | Amp$^r$; *E. coli* pMA-RQ SEanti2 shuttle *Hv*Fdx CRISPRi spacer sequence 1 | This study |
| pJAM4486 | Amp$^r$; *E. coli* pMA-RQ SEanti2 shuttle *Hv*Fdx CRISPRi spacer sequence 2 | This study |
| pJAM4487 | Amp$^r$; *E. coli* pMA-RQ SEanti2 shuttle *Hv*Fdx CRISPRi spacer sequence 3 | This study |
| pJAM4488 | Amp$^r$; *E. coli* pMA-RQ SEanti2 shuttle *Hv*Fdx CRISPRi spacer sequence 3 | This study |
| pJAM4489 | Amp$^r$; Nov$^r$; *H. volcanii*-*E. coli* pJAM4484 shuttle *Hv*Fdx CRISPRi spacer sequence 1 | This study |
| pJAM4490 | Amp$^r$; Nov$^r$; *H. volcanii*-*E. coli* pJAM4484 shuttle *Hv*Fdx CRISPRi spacer sequence 2 | This study |
| pJAM4491 | Amp$^r$; Nov$^r$; *H. volcanii*-*E. coli* pJAM4484 shuttle *Hv*Fdx CRISPRi spacer sequence 3 | This study |
| pJAM4492 | Amp$^r$; Nov$^r$; *H. volcanii*-*E. coli* pJAM4484 shuttle *Hv*Fdx CRISPRi spacer sequence 4 | This study |
| pJAM4547 | Amp$^r$; *H. volcanii*-*E. coli* shuttle P$_{tnaA}$ *Hv*Fdx-His$_6$ K97R expression | This study |
| pJAM4548 | Amp$^r$; *H. volcanii*-*E. coli* shuttle P$_{tnaA}$ *Hv*Fdx-His$_6$ K97Q expression | This study |
| pJAM4549 | Amp$^r$; *H. volcanii*-*E. coli* shuttle P$_{tnaA}$ *Hv*Fdx-His$_6$ K113R expression | This study |
| pJAM4550 | Amp$^r$; *H. volcanii*-*E. coli* shuttle P$_{tnaA}$ *Hv*Fdx-His$_6$ K113Q expression | This study |
| pJAM4551 | Amp$^r$; *H. volcanii*-*E. coli* shuttle P$_{tnaA}$ *Hv*Fdx-His$_6$ K119R expression | This study |
| pJAM4552 | Amp$^r$; *H. volcanii*-*E. coli* shuttle P$_{tnaA}$ *Hv*Fdx-His$_6$ K119Q expression | This study |
| **HVO_2874 (GNAT homolog)-derived** | | |
| pJAM4473 | Amp$^r$; pTA131-based *hvo_2874* pre-deletion plasmid | This study |
| pJAM4474 | Amp$^r$; pTA131-based Δ*hvo_2874* pre-deletion plasmid | This study |
