## Supplemental Table S4 for "Lysine acetylation-mediated regulation of ferredoxin and ferredoxin reductase redox-active proteins in *Haloferax volcanii*"

**Supplemental Table S4.** List of primers used to generate gene deletions, expression plasmids, and site-directed mutagenesis (SDMs).

| Primer | Sequence (5' to 3') | Source |
|---|---|---|
| **HVO_2345 FdR** | | |
| FdR SDM Anchor F1 | GAGATTCTCGGCGGCACCGACATCGAG | This study |
| FdR K378R R1 | GAGCTTCTTGTA**cct**CGTCTGCGGCGAG | This study |
| FdR K378Q R1 | GAGCTTCTTGTA**ctg**CGTCTGCGGCGAG | This study |
| FdR K360R R1 | CGACGATTTTGCCGTC**gcg**GATGGCGAGACGCCGC | This study |
| FdR K360Q R1 | CGACGATTTTGCCGTC**ctg**GATGGCGAGACGCCGC | This study |
| FdR K307R R1 | GTAGTCGACCATGTT**cct**GGCCGCGATGG | This study |
| FdR K307Q R1 | GTAGTCGACCATGTT**ctg**GGCCGCGATGG | This study |
| FdR SDM Anchor R2 | GGAGCCCCACGCGCCGTTCTGGGCGCGG | This study |
| FdR K137R F2 | GCCCGCGACATC**cgc**GAGCACATCGAG | This study |
| FdR K137Q F2 | GCCCGCGACATC**cag**GAGCACATCGAG | This study |
| FdR K176R F2 | CTCATGCGCGGC**cgc**GCGTGGTGGCGC | This study |
| FdR K176Q F2 | CTCATGCGCGGC**cag**GCGTGGTGGCGC | This study |
| **HVO_2995 Fdx** | | |
| Fdx-His$_6$ - pET24b NdeI | TTGCATATGCCCACGGTAACCTACCTC | This study |
| Fdx-His$_6$ - pET24b XhoI | TTTCTCGAGGATGACGCGGTTCTGC | This study |
| Fdx-His$_6$ - pJAM809 NdeI | TTGCATATGCCCACGGTAACCTACCTC | This study |
| Fdx-His$_6$ - pJAM809 KpnI | TTAGGTACCGATGACGCGGTTCTGC | This study |
| Fdx-His$_6$ - pJAM202 NdeI | TTGCATATGCCCACGGTAACCTACCTC | This study |
| Fdx-His$_6$ - pJAM202 BlpI | TAAGCTCAGCGGTGGCAGCAGCCAACT | This study |
| Fdx-His$_6$ - pTA963 NdeI | TTGCATATGCCCACGGTAACCTACCTC | This study |
| Fdx-His$_6$ - pTA963 NotI | TAAGCGGCCGCTGGCAGCAGCCAACTC | This study |
| Fdx SDM Anchor F1 | GTGAGATGGAAGTGAACCAGGGCGAGTAC | (80) |
| Fdx K97Q R1 | GTGAGGCGGACGTT**ctg**CTCGTTGACTTCC | (80) |
| Fdx K97R R1 | GTGAGGCGGACGTT**ccg**CTCGTTGACTTCC | (80) |
| Fdx K113Q R1 | GTTGTAGACGAT**ctg**GACCTCGTCCTC | This study |
| Fdx K113R R1 | GTTGTAGACGAT**ccg**GACCTCGTCCTC | This study |
| Fdx K119Q R1 | GTAGTCGAGGTG**ctg**CGCGTTGTAGAC | (80) |
| Fdx K119R R1 | GTAGTCGAGGTG**ccg**CGCGTTGTAGAC | (80) |
| pJAM202 inverse CRISPR KpnI F | TAAGGTACCCTAGAAGCTTGGATCGGGGGTTGTT | This study |
| pJAM202 inverse CRISPR BamHI R | TAAGGATCCCTGAAAGGAGGAACTATATCCGGATTGG | This study |
| pMA-PQ tele RNase anti 2 KpnI F | TAAGGTACCGAGAATCGAAACGC | This study |
| pMA-PQ tele RNase anti 2 BamHI R | TAAGGATCCCAAAAAACCCCTCAA | This study |
| CRISPR pJAM check F | GCGGTCGGACAACAACCCCCGATCCAA | This study |
| CRISPR pJAM check R | GGGAAAGCCGGCGAACGTGGCG | This study |
| pMA-RQ SEanti2 Check F | GACGGCCAGTGAGCGCGACGTAATAC | This study |
| pMA-RQ SEanti2 Check R | GCGTCGATTTTTGTGATGCTCGTC | This study |

| | | |
|---|---|---|
| Fdx CRISPRi spacer sequence 1 F | CTCTGTCGATGCCCACGGACCGATATTGGTATGGCA | This study |
| Fdx CRISPRi spacer sequence 1 R | GATTCGAAGGCCACCCCAGCTTCAACTACCGATCAA | This study |
| Fdx CRISPRi spacer sequence 2 F | TTCTCGACGACAACGGCTACCGATATTGGTATGGCA | This study |
| Fdx CRISPRi spacer sequence 2 R | CTTCGTAGTTGAGGTAGGGCTTCAACTACCGATCAA | This study |
| Fdx CRISPRi spacer sequence 3 F | CACGGTAACCTACCTCAAACCGATATTGGTATGGCA | This study |
| Fdx CRISPRi spacer sequence 3 R | GGCATCGACAGAGGATTCGCTTCAACTACCGATCAA | This study |
| Fdx CRISPRi spacer sequence 4 F | GTCTAATGGGGTGGCCTTACCGATATTGGTATGGCA | This study |
| Fdx CRISPRi spacer sequence 4 R | TTTGCTGTTCCGTGCGCCGCTTCAACTACCGATCAA | This study |
| **HVO_2874** | | |
| HVO_2874 knock-in BamHI | TAAGGATCCGAATCGTCTCGGTGAGCCGCTGGGTG | This study |
| HVO_2874 knock-in HindIII | TTTAAGCTTGGTTCGGCATGTGGTGGAAGGCGTCC | This study |
| HVO_2874 inverse knock-out F | AACCCTCGGCCCCCGAAAG | This study |
| HVO_2874 inverse knock-out R | GGCCGAACTGGGGCCGCCGCG | This study |
| **qRT-PCR primers** | | |
| RpL-16 (HVO_0484) F | GCGAGTACATCACGGGTATC | (99) |
| RpL-16 (HVO_0484) R | CACTTCCTCTTCGACCTTCAG | (99) |
| Fdx (HVO_2995) F | CCCACGGTAACCTACCTCAAC | This study |
| Fdx (HVO_2995) R | GGATGTACTCGCCCTGGTTC | This study |

Primer sequence: underlined nucleotides represent restriction modification sites and lowercase nucleotides represent site-directed mutations.
