## Supplemental Figures S1-S4 for "Lysine acetylation-mediated regulation of ferredoxin and ferredoxin reductase redox-active proteins in *Haloferax volcanii*"

**A**

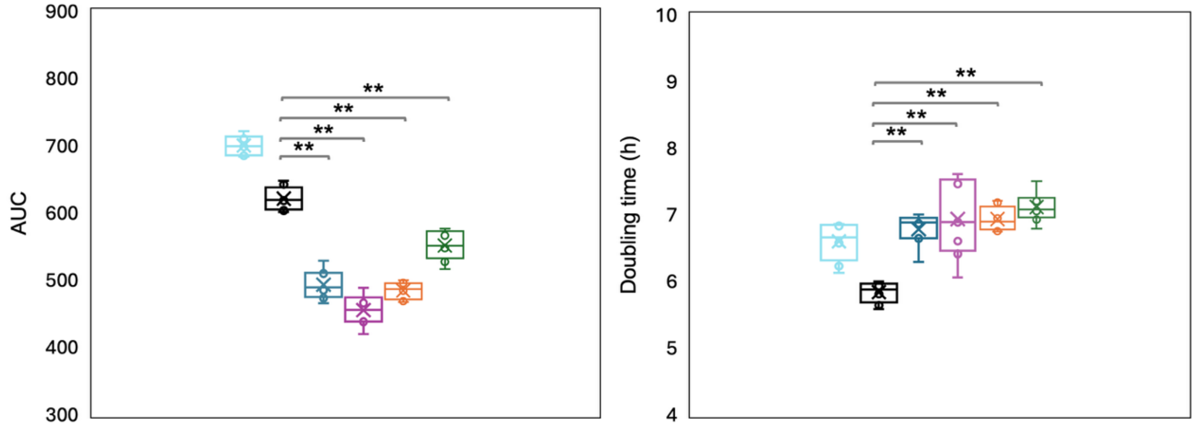


**B**

| Strain | AUC | Growth rate ($h^{-1}$) | Doubling time (h) | Max $OD_{600}$ |
|---|---|---|---|---|
| **HV30** | 700 ± 14 | 0.11 | 6.60 ± 0.28 | 0.50 ± 0.01 |
| **Empty vector** | 621 ± 17 | 0.12 | 5.84 ± 0.15 | 0.46 ± 0.01 |
| **CRISPR *fdx* 1** | 494 ± 22 | 0.10 | 6.79 ± 0.24 | 0.40 ± 0.02 |
| **CRISPR *fdx* 2** | 456 ± 23 | 0.10 ± 0.01 | 6.94 ± 0.57 | 0.38 ± 0.01 |
| **CRISPR *fdx* 3** | 485 ± 12 | 0.10 | 6.94 ± 0.18 | 0.39 ± 0.01 |
| **CRISPR *fdx* 4** | 552 ± 22 | 0.10 | 7.12 ± 0.22 | 0.43 ± 0.10 |

**Supplemental Figure S1.** CRISPRi mediated repression of *fdx* (*ferA5*).

A. *H. volcanii* strains included: HV30 (parent, light blue), HV30/pJAM4484 (empty vector, black), HV30/pJAM4489 (CRISPRi plasmid *fdx* 1, dark blue), HV30/pJAM4490 (CRISPRi *fdx* plasmid 2, purple), HV30/pJAM4491 (CRISPRi *fdx* plasmid 3, orange), HV30/pJAM4491 (CRISPRi *fdx* plasmid 4, green). Growth curve at 42 °C in ATCC974 rich medium supplemented with novobiocin for plasmid selection. Growth was then monitored by $OD_{600}$ using EPOCH2 microplate reader and Gen5 software, every 15 min for 99 h, shaking at double orbital (continuously) (see Methods for details). A student's t-test was used to determine the statistical significance (*p*-value <0.005, **; ns, not significant) of the area under the curve (AUC; **left**) and doubling time ($t_d$; **right**) calculated over the 42 h time course for EV compared to CRISPRi *fdx* plasmid 1 ($p = 5.64 \times 10^{-9}$ for AUC and $7.27 \times 10^{-7}$ for $t_d$), CRISPRi *fdx* plasmid 2 ($p = 6.81 \times 10^{-10}$ for AUC and $7.68 \times 10^{-4}$ for $t_d$), CRISPRi *fdx* plasmid 3 ($p = 1.33 \times 10^{-10}$ for AUC and $3.44 \times 10^{-9}$ for $t_d$), and CRISPRi *fdx* plasmid 4 ($p = 6.56 \times 10^{-6}$ for AUC and $7.00 \times 10^{-9}$ for $t_d$).

B. Growth parameters of CRISPR *fdx* repression. Area Under the Curve (AUC), growth rates, and doubling times of parent (H1207), mutant strain, and complementation cultivated in ATCC974. AUC, area under the curve calculated from 0 to 42 h; growth rate and doubling time determined from 12 h to 24 h; maximum $OD_{600}$ based on $OD_{600}$ reached at 42 h when grown in a 96-well plate.

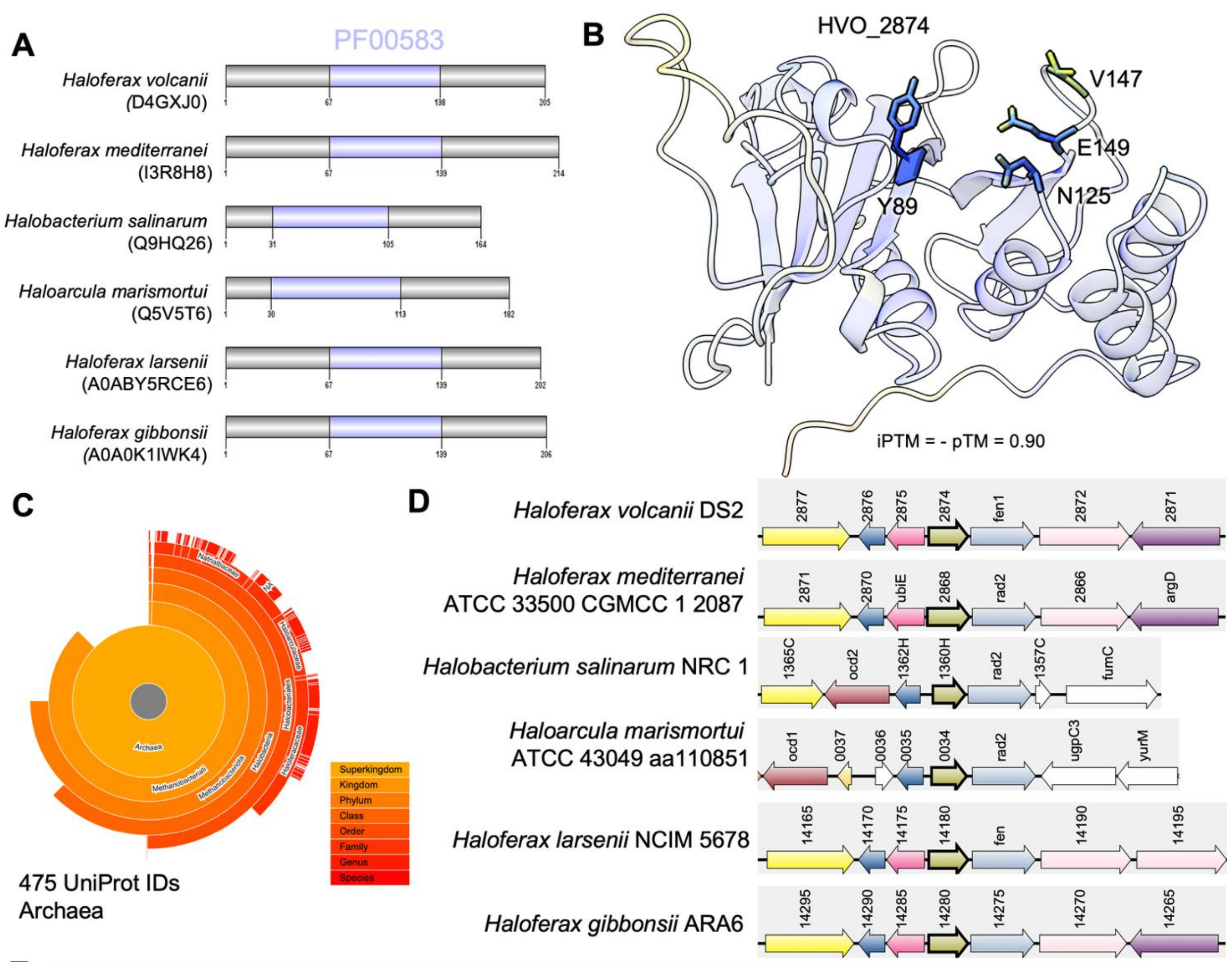


E

| Gene locus tag | Pfam | Description |
|---|---|---|
| HVO_2872 | PF26790<br>PF26793 | Twin arginine translocation (TAT) motif protein |
| HVO_2873 | PF00867<br>PF00752 | Flap endonuclease 1 |
| HVO_2874 | PF00583 | Putative acetyltransferase |
| HVO_2875 | PF08241 | S-adenosylmethionine-dependent methyltransferase |
| HVO_2876 | PF11255 | Uncharacterized, DUF3054 |
| HVO_2877 | PF07690 | Major facilitator superfamily transport protein |

**Supplemental Figure S2.** Structural, phylogenetic, and genomic analysis of the putative acetyltransferase *hvo*_2874.

A. Domain organization of *hvo_2874* homologs identified across haloarchaeal species. Conserved GNAT-family acetyltransferase domains (PF00583) are highlighted in purple, demonstrating conservation of the predicted catalytic acetyltransferase region .
B. AlphaFold structural model of HVO_2874 highlighting residues predicted to participate in acetyl-CoA binding (Y89, N125, V147) and catalysis E149). Structural modeling predicted E149 proximal to *Hv*Fdx K119, supporting a potential role in enzymatic lysine acetylation.
C. Phylogenetic distribution of the top 1000 protein homologs identified using *Hv*HVO_2874 protein sequence as the input for Blast and EFI-EST default parameters. EFI-EST analysis revealed that HVO_2874 homologs were detected exclusively within archaeal lineages. Conserved genomic organization of hvo_2874 homologs among 6 representative haloarchaeal species. Arrows represent open reading frames (ORFs) with gene locus tag numbers.
D. Genomic neighborhood of *Haloferax volcanii hvo_2874*. Pfam, database of protein family annotations.
E. Gene locus tag, Pfam and description of *hvo_2874* genomic neighborhood.

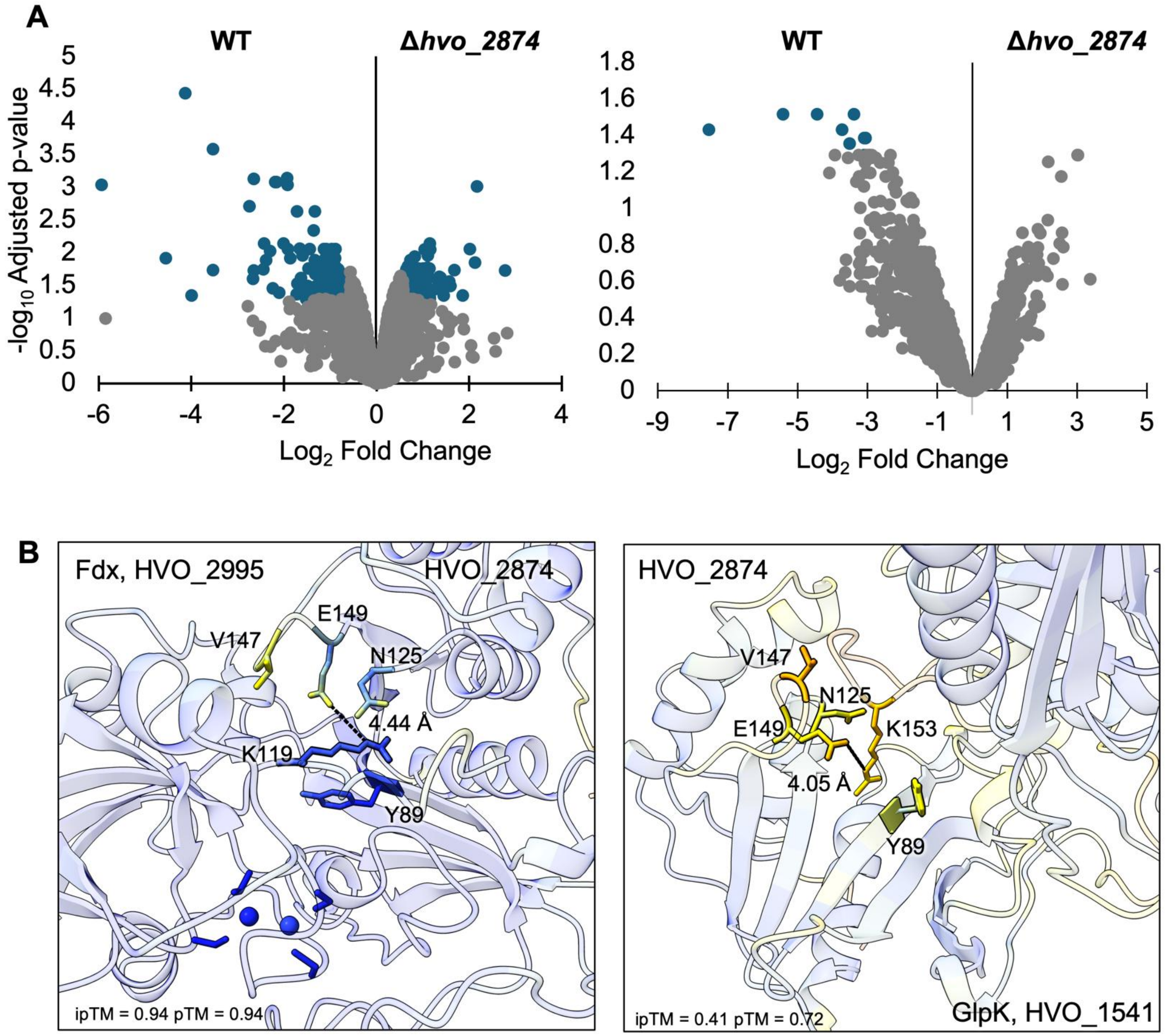


**Supplemental Figure S3.** Deletion of *hvo_2874* alters protein abundance and lysine acetylation profile in *H. volcanii*.

A. Differential protein abundance profiles associated with deletion of *hvo_2874*. Volcano plots showing significantly altered proteins grown in 'SLG' (left) and glycerol (right) in the Δ*hvo_2874* strain relative to the parental strain. Data are displayed as $\log_2$ fold change versus $-\log_{10}$ adjusted p-value. Significantly enriched features are highlighted in blue, while non-significant features are shown in gray.

B. AlphaFold structural predictions were used to model interactions between the lysine acetyltransferase HVO_2874 and two putative substrate proteins identified from acetylome profiling: HVO_2995 (Fdx, K119) and HVO_1541 (GlpK, K153). The following was also detected in the acetylome profile, though at large spatial separations: HVO_0769 TRAM domain protein (21.13 Å), HVO_1797 beta-lactamase domain protein (46.72 Å), and HVO_A0551 (Acs9) acetyl-CoA synthetase (31.49 Å). The active site of HVO_2874, centered on glutamate 149 (E149), and surrounding putative acetyl-CoA–binding residues (Val147, Asn125, Tyr89) are indicated. Acetyl-lysines, catalytic residue E105, and distances (Å) are indicated.

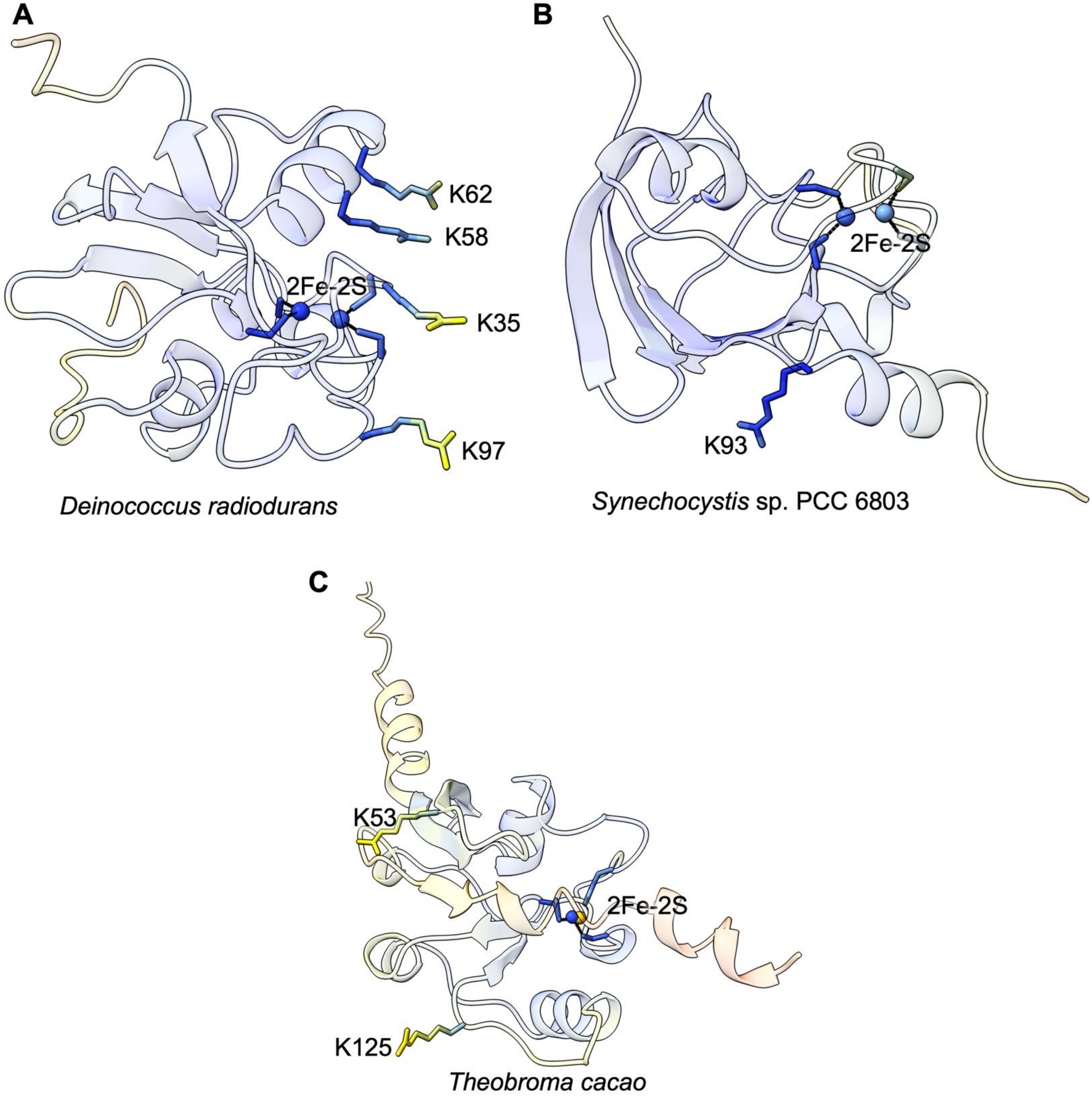


**Supplemental Figure S4.** Comparison of lysine-acetylated ferredoxins from diverse organisms. Structural models of ferredoxins reported to undergo lysine acetylation in other organisms are shown with acetylated lysine residues highlighted and the 2Fe–2S cluster indicated. The upper left (A) panel shows a ferredoxin from *Deinococcus radiodurans* (DR_2075) containing four acetylated lysine residues (K35, K58, K62, and K97) near the iron sulfur cluster. The upper right (B) panel displays the ferredoxin from *Synechocystis* sp. PCC 6803 (Slr1828) with a single acetylated residue (K93), and the lower (C) panel displays a ferredoxin with acetylation at K53 and K125 from *Theobroma cacao* (TCM_002657). Among the acetylation sites identified across these ferredoxins, only K35 and K97 from *D. radiodurans* is positioned in close proximity to the 2Fe–2S cluster, whereas the remaining acetylated lysine residues are located distal to the 2Fe-2S cluster.
